# Map of epigenetic age acceleration: a worldwide meta-analysis

**DOI:** 10.1101/2024.03.17.585398

**Authors:** Igor Yusipov, Alena Kalyakulina, Claudio Franceschi, Mikhail Ivanchenko

## Abstract

This study is the first systematic meta-analysis of epigenetic age acceleration of the largest publicly available DNA methylation data for healthy samples (93 datasets, 23K samples), focusing on geographic and ethnic aspects of different countries (25 countries) and populations (31 ethnicities) around the world. The most popular epigenetic tools for assessing age acceleration were examined in detail, their quality metrics were analyzed, and their ability to extrapolate to epigenetic data from different tissue types and age ranges different from the training data of these models was explored. In most cases, the models are not consistent with each other and show different signs of age acceleration, with the PhenoAge model tending to systematically underestimate and different versions of the GrimAge model tending to systematically overestimate the age prediction of healthy subjects. Although GEO is the largest open-access epigenetic database, most countries and populations are not represented, and different datasets use different criteria for determining healthy controls. Because of this, it is difficult to fully isolate the contribution of “geography/environment”, “ethnicity” and “healthiness” to epigenetic age acceleration. However, the DunedinPACE metric, which measures aging rate, adequately reflects the standard of living and socioeconomic indicators in countries, although it can be applied only to blood methylation data. When comparing epigenetic age acceleration, males age faster than females in most of the countries and populations considered.

## 1. Introduction

Aging studies are recently one of the most important research areas, where accurate measurement of biological aging, prediction and identification of risks of age-associated diseases and mortality are very important parts [1], as well as the discovery of both effective preventive health promotion methods and interventions aimed at modulating the aging process [2]. The aging process is associated with changes at many levels of human body functioning, among which an important element is epigenetic changes - genome modifications associated not with DNA sequence changes but with chromatin modifications [3]. The release of high-throughput arrays measuring DNA methylation [4–6] has stimulated research on epigenetic changes in the context of biological age estimation [7,8], the relationship between epigenetic age and age-associated pathologies [9], and the ability to assess the efficacy of anti-aging interventions [10,11]. Such studies strictly separate chronological age and biological age. Chronological age usually refers to the actual years of a person’s life, while biological age refers to the physiological condition and functioning of the human body [9]. A person may be older or younger by biological age than by chronological age, which reflects the person’s state of health and aging and provides a relative measure of how well the person’s body is functioning compared to that person’s chronological age [12].

Various biomarkers, including epigenetic ones, can be used to estimate biological age. The most common is the estimation of DNA methylation age using epigenetic clock models [13]. The best known epigenetic clocks are Horvath’s clocks [14], which can estimate the age of many tissues (so-called pan-tissue clocks). Another Horvath’s clock can work more accurately for cultured cells, such as fibroblasts [15]. About the same time, no less famous Hannum clocks [16] were developed for blood DNA methylation (so-called tissue-specific or single-tissue clocks). The first generation of epigenetic clocks was not sufficiently correlated with clinical blood measures, and to include these biomarkers the DNAm PhenoAge epigenetic clock was developed [17], regressing these blood measures on DNA methylation data and then estimating age. DNAm PhenoAge can differentiate morbidity and mortality risks in people of the same chronological age [17]. The GrimAge clock is composed of seven plasma protein markers based on DNA methylation and the number of smoking years that are associated with morbidity or mortality [18]. Compared to other models, GrimAge focuses more on lifestyle and age-related conditions, so it predicts longevity well [18]. Another epigenetic metric often considered together with the epigenetic clocks is the DunedinPACE aging rate [19], which corresponds to the number of biologic years per chronological year. This metric is associated with morbidity, disability, and mortality. However, different epigenetic clocks/metrics show varying levels of association with risk factors and the consequences of aging [8], suggesting that they may also reflect distinct aspects of aging [20].

Often, epigenome studies do not emphasize the racial/ethnic composition of the data, but many results are based on Caucasians and cannot always be generalized to other groups [21,22]. Racial/ethnic and socio-demographic information is important precisely to assess whether epigenetic clocks can be generalized to different groups of people, given that variations in DNA methylation are altered by genetic, social, and environmental factors [23–27], and unfavorable social and biophysical factors can influence disease risk and healthy aging processes [28,29]. This is a known problem when prediction models are ineffective in social groups that were under/unrepresented in the training data used to build the models [30,31], and estimates will then be less accurate [32]. One of the directions for future experiments and recommendations in [3] is the detailed analysis of different populations around the world.

To date, there is no systematic meta-analysis of epigenetic age acceleration in different world regions and populations. In one of the most famous works investigating the running of epigenetic clocks in different races/ethnicities [33], it was found that the rate of epigenetic aging is significantly related to race/ethnicity. A wide range of races were considered (African Ancestry, Caucasians, Hispanics, East Asians, Tsimane Amerindians), but epigenetic markers were limited to IEAA and EEAA based on the Horvath’s model. In a study of the U.S. adult population [34], first-generation epigenetic clocks (Horvath and Hannum) show slower aging for Blacks compared to Whites, second-generation clocks (PhenoAge and GrimAge) and DunedinPACE show faster aging for Blacks, and Hispanics in general are aging more slowly. The influence of race on epigenetic age acceleration is also explored in papers [35–39], demonstrating the importance of addressing the racial/ethnic component in DNA methylation data during research. Thus, the existing works miss one or more of the following important points: (i) they consider a limited number of populations (often within the same region); (ii) they do not take into account the complexities caused by batch effects: methylation values strongly depend on the quality of the raw data and its preprocessing, very different results can be obtained in different laboratories; (iii) they use only the first-generation, most common versions of epigenetic clocks (Horvath, Hannum), without taking into account more modern epigenetic metrics.

In this paper, we performed a systematic meta-analysis and studied epigenetic age acceleration worldwide using the largest DNA methylation datasets from GEO [40], the largest open access repository. We investigated how healthy individuals from different countries and populations (we took only healthy controls) differ in the most well-known epigenetic metrics (Horvath DNAmAge [14], Hannum DNAmAge [16], SkinBlood DNAmAge [15], DNAm PhenoAge [17], GrimAge [18], their PC-modifications [41], GrimAge version 2 [42], DunedinPACE [19]) and analyzed their quality metrics, possibilities to extrapolate to epigenetic data from different tissue types and age ranges different from the training data of these models. In the paper, for all used data, a harmonization procedure was performed for mandatory leveling of batch effects for methodologically correct comparison of datasets. We also addressed epigenetic age acceleration between males and females.

The paper is organized as follows. Section 2.1 describes the analyzed DNA methylation data, the principles of its inclusion/exclusion from the analysis, and the considered epigenetic metrics. Section 2.2 summarizes the results of the analysis of epigenetic age acceleration in different tissues. Section 2.3 compares individuals from different countries and populations in the context of their epigenetic characteristics. Section 2.4 provides more detailed results for individual (most representative) countries.

## 2. Results

### 2.1. Epigenetic data and study design

Figure 1 shows the overall study design, which includes GEO database parsing, preprocessing and harmonization of DNA methylation data, calculation of epigenetic age acceleration for the analyzed epigenetic clock models, and the meta-analysis of epigenetic age acceleration itself in different tissues, countries and populations, sex-specificity is also taken into account. Let us consider each step in more detail.

#### 2.1.1. GEO parsing

We parsed the entire GEO database to select DNA methylation datasets suitable for analysis. We considered the Infinium HumanMethylation450 (Illumina 450k, GEO has 2 codes for this standard, GPL13534 and GPL16304) and Infinium MethylationEPIC (Illumina EPIC, GEO has 2 codes for this standard, GPL21145 and GPL23976) standards. Next, we briefly describe the rules for selecting datasets from GEO for our meta-analysis (Figure 1, Step 1).

**Figure 1.**
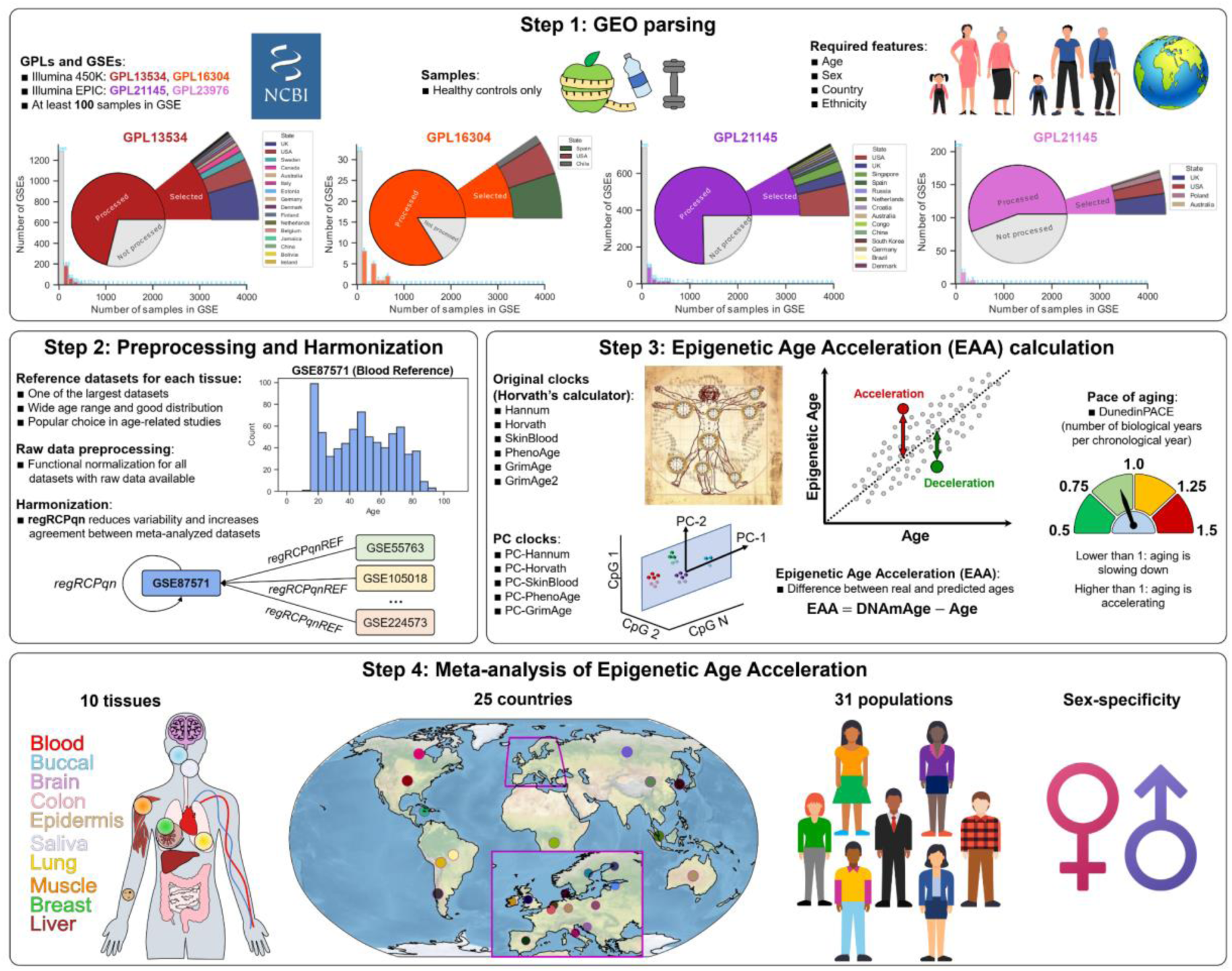
Main steps and design of the study. Step 1: GEO datasets selection. The Illumina 450k and Illumina EPIC standards were considered. For all GPLs representing these standards, the total number of datasets by the number of samples in them is plotted, the proportion of processed and unprocessed datasets, the proportion of datasets selected for meta-analysis, and their representation by country are shown. Main inclusion criteria: healthy controls; information on age, sex, country of residence/origin, and race/ethnicity is available. The inclusion/exclusion criteria of the datasets are described in more details in Sections 2.1.1, 4.1. Step 2: To correctly perform the meta-analysis, DNA methylation data was harmonized for all used datasets. For each tissue, a reference dataset (usually one of the largest with a wide age range) was selected, relative to which harmonization was performed for all other datasets using the regRCPqnREF approach. Step 3: Reviewed epigenetic metrics (classical epigenetic clock models, their PC-modifications and DunedinPACE aging rate) and the method of epigenetic age acceleration calculation as a machine learning model error. Step 4: Entities for meta-analysis: 10 tissues, 25 countries, 31 populations, sex-specificity.

We select only healthy controls (understanding that the control group may be defined differently across studies), all samples must have sex and age information, and a precise indication of race/ethnicity. We used Geo-referencing Ethnic Power Relations (GeoEPR 2021) to systemize countries and population groups in them [43]. We excluded samples with unknown race/ethnicity and samples with unknown country of origin/residence. Participants must be at least 3 years old. Despite the fact that children for most of the epigenetic clock models were not part of the training data, we believe that it would be interesting to study and analyze the ability of these models to make extrapolated predictions on participants of small age. The dataset must have a release date on GEO no later than December 14, 2023 and initially contain at least 100 individuals (there may be fewer healthy controls). Detailed sample inclusion/exclusion criteria are provided in Methods, Section 4.1.

Figure 1, Step 1 summarizes the statistics of the number of datasets and samples for each GPL standard. For GPL13534 (Illumina 450k), 334 datasets were processed, containing 102358 samples; 56 datasets were selected for meta-analysis, containing 15549 samples. For GPL16304 (Illumina 450k doublet, contains significantly fewer datasets), 17 datasets were processed, containing 5129 samples; 3 datasets were selected for meta-analysis, containing 591 samples. For GPL21145 (Illumina EPIC), 174 datasets were processed, containing 57913 samples; 30 datasets were selected for meta-analysis, containing 6309 samples. For GPL23976 (Illumina EPIC doublet, contains significantly fewer datasets), 29 datasets were processed, containing 8903 samples; 4 datasets were selected for meta-analysis, containing 767 samples. All 93 selected datasets are listed in the Data Availability Statement. Detailed information about all processed datasets (number of samples, available characteristics) with a description of the reasons for their inclusion or exclusion from analysis is given in Supplementary Tables S1-S4.

#### 2.1.2. Preprocessing and Harmonization

DNA methylation datasets in GEO can be divided into 2 large groups by storing type: datasets with raw data (idat files that can be processed in any chosen way) and datasets with already preprocessed data (can be preprocessed in many ways and presented in different resulting formats). Different types of preprocessing are one of the sources of batch effects (along with different laboratory conditions and many other factors) that can influence the results of meta-analyses and introduce non-biological sources of variability. It has been shown in [44] that results can vary greatly depending on the analyzed dataset, and data harmonization is one of the key steps in meta-analysis. For harmonization of DNA methylation data used in this paper, we used the regRCPqnREF (regional Regression on Correlated Probes with quantile normalization and reference) approach, which harmonizes multiple DNA methylation datasets relative to some selected reference dataset (usually a rather large by number of samples dataset with a wide age range and close to uniform distribution of samples across age groups). For each considered tissue type, a different reference dataset was selected (in case more than 1 dataset is available for a given tissue type). For DNA methylation in blood, the reference dataset was GSE87571 (Figure 1, Step 2), for buccal cells - GSE137688, for brain - GSE74193, for colon - GSE151732, for saliva - GSE111223.

#### 2.1.3. Epigenetic Age Acceleration

Table 1 summarizes the main characteristics of the epigenetic models used in this paper. Most of the considered epigenetic clock models are single-tissue epigenetic age estimates that were built only on blood data. The DunedinPACE aging rate is also a single-tissue metric. Only two Horvath epigenetic clock models (Horvath DNAmAge and SkinBlood DNAmAge) with their PC modifications are pan-tissue models built on data from multiple heterogeneous tissues. Such models have higher applicability to a wide range of data, can estimate separately the aging rate of different organs in the human body, and are potentially more sensitive to different pathological conditions. The data on which the considered models were trained are also significantly different - most epigenetic clocks were built on a few closed-access datasets, however, rather large in the number of samples. Interestingly, almost all epigenetic models were either built entirely on data from US representatives or have datasets with US representatives among the training datasets. When applying these clock models to different population cohorts, their age acceleration will be considered relative to exactly the training datasets of the clock models, which are mostly US. The relatively simple linear ElasticNet model was used to construct all the considered epigenetic metrics models (the best known and most used models).

**Table 1.** Epigenetic clock models analyzed in this paper. For each model, the following information is given: method of construction, number of training datasets, number of samples in training datasets, representatives of which countries are in the training datasets, whether the training data are open or closed access, whether the model is single-tissue or pan-tissue.

| Epigenetic clock | Method | Number of train datasets | Number of samples in train datasets | Train data origin | Open or closed train data | Single-tissue or pan-tissue |
| --- | --- | --- | --- | --- | --- | --- |
| Hannum DNAmAge | ElasticNet | 1 dataset | 482 | USA | Open | Single-tissue (blood) |
| Horvath DNAmAge | ElasticNet | 39 datasets | 3931 | Multiple countries | Partially open | Pan-tissue |
| SkinBlood DNAmAge | ElasticNet | 10 datasets | 896 | Multiple countries | Partially open | Pan-tissue |
| DNAm PhenoAge | Cox regression + ElasticNet | 2 datasets | 9926 (step 1)<br>912 (step 2) | USA, Italy | Closed | Single-tissue (blood) |
| GrimAge | ElasticNet + Cox regression | 1 dataset | 1731 | USA | Closed | Single-tissue (blood) |
| GrimAge version 2 | ElasticNet + Cox regression | 1 dataset | 1833 | USA | Closed | Single-tissue (blood) |
| PC-Hannum | PCA + ElasticNet | 1 dataset | 656 | USA | Open | Single-tissue (blood) |
| PC-Horvath | PCA + ElasticNet | 37 datasets | 4297 | Multiple countries | Partially open | Pan-tissue |
| PC-SkinBlood | PCA + ElasticNet | 9 datasets | 895 | Multiple countries | Open | Pan-tissue |
| PC-PhenoAge | PCA + ElasticNet | 2 datasets | 4505 | USA, Italy | Closed | Single-tissue (blood) |
| PC-GrimAge | PCA + ElasticNet | 2 datasets | 2754 | USA | Closed | Single-tissue (blood) |
| DunedinPACE | Linear mixed-effects model + ElasticNet | 1 dataset | 1037 | New Zealand | Closed | Single-tissue (blood) |

All epigenetic clocks are machine learning regression models, and their performance should be analyzed using conventional machine learning metrics. Often age acceleration is defined as residuals relative to a linear regression built on some subset [21,34,45,46] because many clock models have systematic bias and their predicted values differ quite significantly from the chronological age (the cloud of points on the Age - Estimated Age plane (Figure 1, Step 3) has a slope different from the bisector of this plane). Such differences may be due to a significant bias between the data on which the clock was trained and the test data on which age acceleration is investigated. This approach to determine age acceleration may be useful in the relative comparison of several different groups [47–49]. From a machine learning perspective, it is methodologically correct to consider the error - the direct difference between predicted and chronological age, so in this study we defined age acceleration in this way. In this paper, we wanted to analyze how all the clock models perform for all the studied populations, as is, with bias and errors. Since we analyzed healthy controls, systematic significant deviations in the clocks will not be due to pathological conditions, but to peculiarities of the clock models themselves.

#### 2.1.4. Meta-analysis of Epigenetic Age Acceleration

Next, we conducted a comparative meta-analysis of epigenetic age acceleration across different countries and ethnic groups, both within and between countries, the results are described in detail in Sections 2.2-2.4. The influence of methylation differences in different tissues of the human body was taken into account, as well as the fact that different clocks have different predictive potential for different age groups. The sex-specific aspect was also addressed (Figure 1, Step 4). Information about all samples involved in the analysis is presented in Supplementary Table S5.

### 2.2. Epigenetic age acceleration in different tissues

There are many studies demonstrating that DNA methylation of different tissues in the human body can vary greatly [50–54], also changing with age [12,55]. Due to this fact, there are different epigenetic clock models - some of them, such as Hannum DNAmAge, DNAm PhenoAge, GrimAge 1 and 2, DunedinPACE, focus only on blood DNA methylation, while others, such as Horvath DNAmAge and SkinBlood DNAmAge are able to analyze the epigenetic age of multiple tissues. Nevertheless, it would be interesting to examine how the considered epigenetic metrics are applied to different human organs and tissues, whether there are any common trends between epigenetic models, how they differ, whether it makes any sense to apply methylation models developed on blood to other tissues.

Figure 2 shows the main results of the analysis of different epigenetic clocks for all the considered tissue groups. The tissues involved in the analysis are shown on the left side in the schematic representation of a human. It is important to note that the analysis considers aggregated tissues groups - all tissue types related to blood (whole blood, peripheral blood, monocytes, etc.) are combined into a single group “Blood”, all tissue types related to the brain (different parts of the brain) are combined into a single group “Brain”, and so on. The resulting number of samples for all 10 tissue types with their corresponding age distributions are shown in Figure 2A. Expectedly, DNA methylation data from blood have the most samples, with one of the widest age distributions. Buccal cells come second, with a high representation of young samples and a lack of samples older than 60 years. The narrowest age distributions are in epidermis (40-80 years) and muscle (20-50 years). Both of these factors, number of samples and age distribution, can influence the performance and quality of epigenetic clocks, as their training data can differ significantly in these two factors.

**Figure 2.**
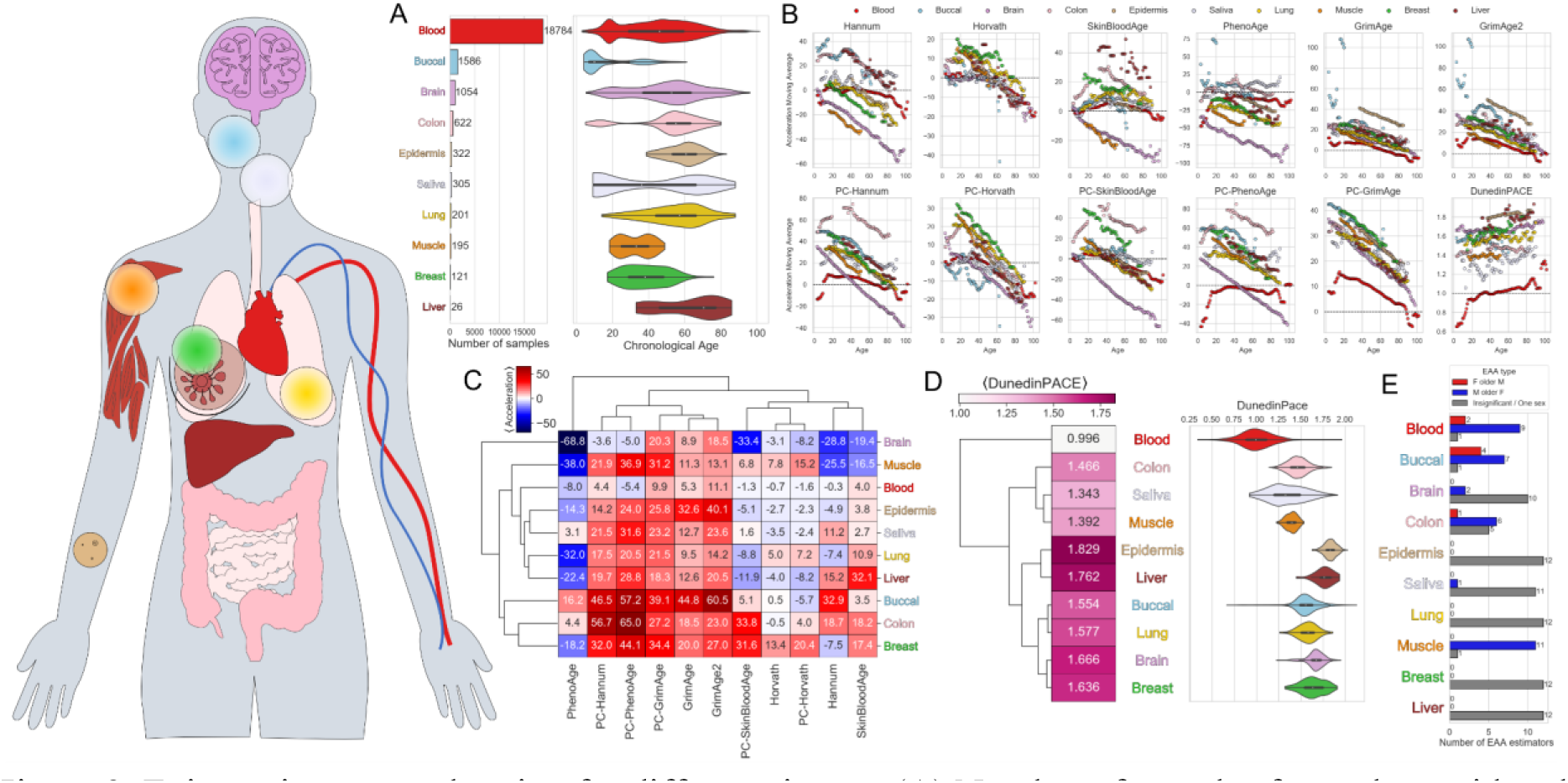
Epigenetic age acceleration for different tissues. (A) Number of samples for each considered tissue and the corresponding age distributions. Each tissue type is matched to a different color. (B) Dependence of the moving average (within 5 years) of epigenetic age acceleration on real age for all 10 tissues for all considered epigenetic models. (C) Diagram of the mean values of epigenetic age acceleration for 10 tissues for all considered epigenetic clocks with hierarchical clustering. Color indicates the value of the mean epigenetic age acceleration. (D) Diagram of mean DunedinPACE epigenetic aging rate values for 10 tissues with hierarchical clustering (left) and corresponding distributions of DunedinPACE values (right). The value of the mean DunedinPACE is indicated in color. (E) Number of epigenetic metrics for each tissue type for which: males have higher epigenetic age acceleration (blue), females have higher epigenetic age acceleration (red), no statistical significance/one sex (gray). Mann-Whitney U-test was applied for males and females for each tissue, with FDR-corrected resulting p-values.

To investigate how the considered epigenetic models work for different tissues at different age ranges, we calculated the moving average (within 5 years) for epigenetic age acceleration values as a function of the real age of the samples (Figure 2B). Expectedly, for most epigenetic models, blood demonstrates adequate epigenetic age acceleration values, with no extreme low or high values. Hannum and Horvath models show underestimation at old age, DNAm PhenoAge model underestimates across the age range, DunedinPACE aging rate shows underestimation at very young age and overestimation at very old age (possibly because such samples were underrepresented in the training data). Epigenetic age acceleration in the brain behaves similarly in the Hannum, Horvath, SkinBlood, PhenoAge models with their PC modifications - it decreases almost monotonically with age to extremely low values in old age. Buccal cells in PhenoAge and both versions of GrimAge for young samples show a very large overestimation (with epigenetic age acceleration values up to 100 years and more). Interestingly, most tissues in the GrimAge models behave in a similar way, monotonically decreasing with age. DunedinPACE demonstrates adequate aging rate values (about 1) only for blood, strongly overestimating values for all other considered tissues. We should note that DunedinPACE values correspond to the number of biological years in one chronological year; values greater than 1 correspond to accelerated aging, less than 1 to decelerated aging.

Next, we looked at the average values of epigenetic age acceleration for different tissues and different epigenetic models as a diagram with hierarchical clustering (Figure 2C). It is consistent that all epigenetic clocks work best for blood, with age accelerations not taking extremely low or high values. This is mainly because most epigenetic models are either based entirely on blood DNA methylation data, or a large part of the training data contains blood DNA methylation data (along with methylation of other tissues). The Horvath and PC-Horvath models perform reasonably well for almost all tissues (except breast) because they were trained on numerous datasets (39) for many different human organs and tissues. SkinBloodAge and PC-SkinBloodAge also perform rather well for a large number of tissues (they were trained on buccal, skin, and blood data) and it is understandable that they perform worse on brain, colon, breast, muscle, and liver. PhenoAge and PC-PhenoAge, GrimAge, GrimAge2 and PC-GrimAge only do well with blood and adapt poorly to other tissues. Interestingly, Hannum does well on a wider range of tissues (given that it was trained on blood data only) - breast, blood, buccal, saliva, lung. PC-Hannum only does well on blood and brain. Hierarchical clustering by epigenetic age combines Horvath and SkinBloodAge with their PC versions and Hannum age into one group, the second group includes all other clocks except PhenoAge, which is separate from all others and differs greatly in results. Hierarchical clustering by tissue shows that similar tissues in terms of epigenetic age acceleration are epidermis and saliva, lung and liver, blood and muscle. Another group contains colon and breast tissues, buccal, while the brain is separated from all other tissues. Most tissue pairs differ statistically significantly in terms of mean epigenetic age acceleration for almost all epigenetic estimates (more details in Supplementary Materials, Supplementary Figures S1, S2).

Since DunedinPACE does not estimate epigenetic age but the aging rate, we discussed it separately (mean values for the considered tissues and distributions of values are shown in Figure 2D). The lowest value is observed for blood, it is closest to 1 (no acceleration or deceleration). This is an expected result since the metric is trained on blood data only. Next in ascending average aging rate is the group of colon, saliva, and muscle (about 1.3-1.4). For all other tissues, the aging rate is above 1.5 (within the DunedinPACE metric, this is as if the tissues aged more than a year and a half in one chronological year). Given these results for different tissues and the fact that DunedinPACE was built on blood data only, we can say that DunedinPACE is relevant only for blood, whereas for other tissues there is a systematic bias. It is also worth noting that the epidermis and liver data are mostly presented for middle-aged and elderly samples, which may also be one of the reasons for the rather large DunedinPACE values.

We also, for each tissue, divided the entire population into females and males, and applied the Mann-Whitney U-test for these two groups [56], obtaining FDR-adjusted p-values. We then calculated the number of epigenetic clocks that have higher epigenetic age acceleration in males than females (blue bars in Figure 2E), higher epigenetic age acceleration in females than males (red bars in Figure 2E), and no statistical significance (gray bars in Figure 2E). Males have higher epigenetic age acceleration than females for blood, buccal cells, colon, saliva, and muscle. Females do not show higher epigenetic age acceleration than males for any tissue. For the remaining tissues, there are no statistical differences between males and females or only one sex is presented in the data.

Since blood is the most representative in sample size, further results in Sections 2.3 and 2.4 will be presented only for blood DNA methylation. Results for other tissues are presented in Supplementary Materials, Supplementary Figures S5-S11.

### 2.3. Epigenetic age acceleration between the countries and populations

DNA methylation is not constant, but dynamically changes over time under the influence of many factors, both internal and external. People living in different parts of the globe may have different DNA methylation (and consequently different rates of epigenetic aging) not only because of different races/ethnicities, but also because of different environmental conditions, living standards, social conditions, access to medical care and many other factors. It may be difficult to separate the influence of all these factors, however, the study of epigenetic age acceleration in different regions of the world is of interest. We examined in detail blood DNA methylation data (as the largest sample size) in the context of differences in epigenetic age acceleration and aging rates between countries (Figure 3) and populations (Figure 4).

**Figure 3.**
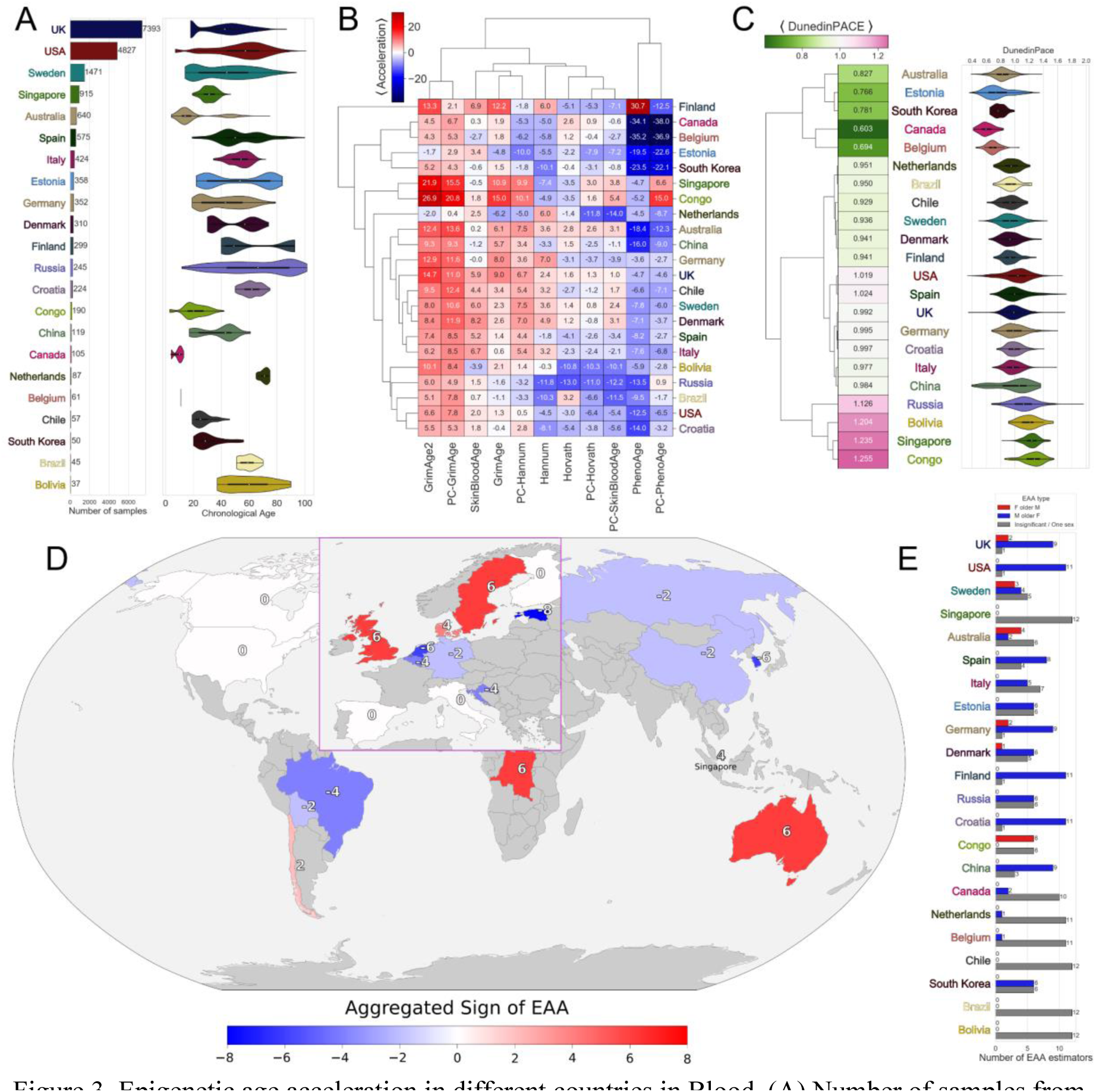
Epigenetic age acceleration in different countries in Blood. (A) Number of samples from different countries (left) with corresponding age distributions (right). (B) Diagram of the mean values of epigenetic age acceleration for countries for all epigenetic clocks with hierarchical clustering. Color indicates the value of the mean epigenetic age acceleration. (C) Diagram of mean DunedinPACE epigenetic aging rate values for all countries with hierarchical clustering (left) and corresponding distributions of DunedinPACE values (right). The value of the mean DunedinPACE is indicated in color. (D) Map showing the Aggregated Sign of <EAA> (difference between epigenetic metrics with positive age acceleration and those with negative age acceleration) for all countries analyzed. Aggregated Sign of <EAA> is also highlighted in color. (E) Number of epigenetic metrics for each country for which: males have higher epigenetic age acceleration (blue), females have higher epigenetic age acceleration (red), no statistical significance/only one sex (gray). Mann-Whitney U-test was applied for males and females for each tissue, with FDR-corrected resulting p-values.

**Figure 4.**
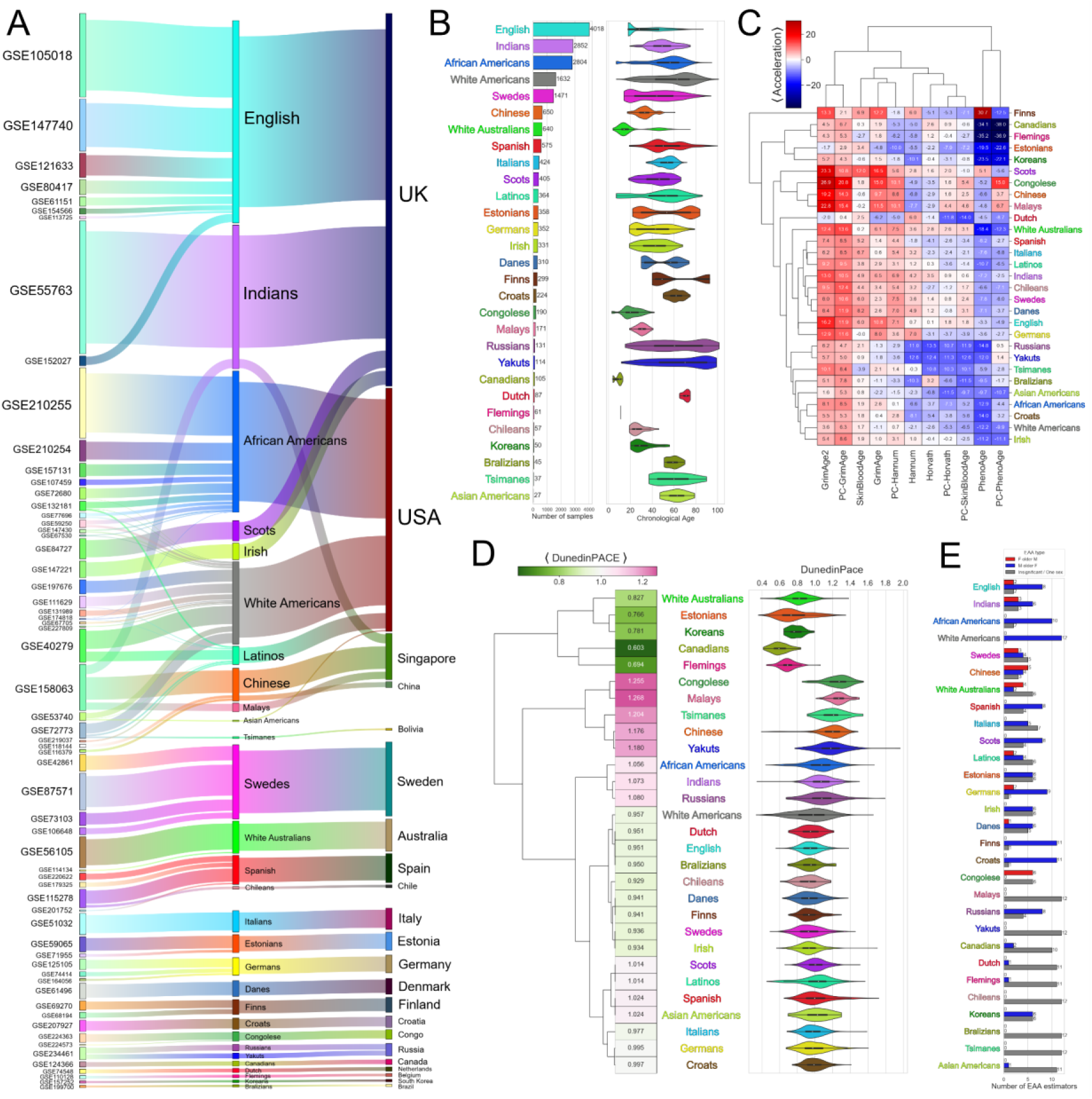
Epigenetic age acceleration in different populations in Blood. (A) Sankey plot showing relations between datasets (GSEs, left), populations (center) and countries (right). (B) Number of samples from different populations (left) with corresponding age distributions (right). (C) Diagram of the mean values of epigenetic age acceleration for populations for all epigenetic clocks with hierarchical clustering. Color indicates the value of the mean epigenetic age acceleration. (D) Diagram of mean DunedinPACE epigenetic aging rate values for all populations with hierarchical clustering (left) and corresponding distributions of DunedinPACE values (right). The value of the mean DunedinPACE is indicated in color. (E) Number of epigenetic metrics for each population for which: males have higher epigenetic age acceleration (blue), females have higher epigenetic age acceleration (red), no statistical significance/only one sex (gray). Mann-Whitney U-test was applied for males and females for each tissue, with FDR-corrected resulting p-values.

The selected datasets of blood DNA methylation show a diversity of representatives from different countries: 22 countries from all continents - Asia, Europe, Africa, America, Australia (Figure 3A). The largest number of samples are from the UK and USA (over 7000 and 4000 samples, respectively). The age range of samples from different countries also differs significantly. The widest age range is observed in the UK, USA, Sweden, Spain, Germany and Russia; samples from Finland are divided into two subgroups with chronological ages of 40-60 years and 80 or more years. Two subgroups are also present in the Estonian data - mostly young 20-40 years and older 60-80 years; in Canada and Belgium all samples are children; the sample in South Korea is mostly young 20-40 years; Singapore and Congo are similar in age range - less than 40 years; samples from the Netherlands are elderly (60-80 years).

The mean values of epigenetic age acceleration for different countries and different epigenetic models are presented as a diagram with hierarchical clustering (Figure 3B). The first observation that stands out is that there is no single country for which all epigenetic clocks give only positive or negative age acceleration (in other words, the results of the models do not agree with each other). PhenoAge and PC-PhenoAge give negative epigenetic acceleration for almost all countries (they are grouped in a separate hierarchical cluster), while GrimAge2 and PC-GrimAge, on the contrary, give positive age acceleration for almost all countries (exactly GrimAge2, not GrimAge) and also form a separate cluster. If we focus our attention on epigenetic acceleration by countries and their hierarchical clustering, we can see that Finland stands apart: PhenoAge, GrimAge and GrimAge2 give large positive age acceleration, PC-PhenoAge - large negative age acceleration (interestingly, there is a huge difference between PhenoAge and PC-PhenoAge), other epigenetic clocks give not very large acceleration of different sign. Next in the hierarchical clustering are Canada and Belgium, Estonia and South Korea, for which PhenoAge and PC-PhenoAge give very large negative age acceleration, and all other clocks show small positive and negative values. Singapore and Congo show large positive GrimAge, GrimAge2 and PC-GrimAge acceleration, and they are almost the only countries with positive acceleration for PC-PhenoAge. For the Netherlands, most of the epigenetic age acceleration rates are negative or slightly positive. Next comes a whole block of similar countries in terms of epigenetic characteristics - Australia, China, Germany, UK, Chile, Sweden, Denmark, Spain, Italy - all of them have strongly positive GrimAge2 and PC-GrimAge acceleration, negative PhenoAge and PC-PhenoAge acceleration. Interestingly, most of the developed European countries with a fairly high quality of life are in this group. The next group includes Bolivia, Russia, Brazil, USA, Croatia - most of them have negative acceleration on Hannum, Horvath, PC-Horvath and PC-SkinBloodAge, rather strongly negative acceleration of PhenoAge, weakly negative PC-PhenoAge. Most country pairs are statistically significantly different in terms of mean epigenetic age acceleration for almost all epigenetic estimates (see Supplementary Materials, Supplementary Figures S3, S4 for details). Since DunedinPACE does not estimate epigenetic age but the aging rate, we discussed it separately (the mean values for the considered countries and the distributions of values are shown in Figure 3C). Canada and Belgium have the lowest aging rates, probably because all the samples are children, and DunedinPACE shows a significant underestimation for children. Australia, Estonia and South Korea are also aging very slowly. Most countries have aging rates close to 1. For them, the DunedinPACE distribution is quite wide, but the mean is close to 1. Samples from Russia, Bolivia, Singapore and Congo age fastest. This seems to be influenced by different factors - quality of life, country development, health care, climate and data particularities. In general, we can conclude that DunedinPACE is a characteristic that is quite strongly related to the quality of life in different countries, showing higher values in developing countries (accelerated aging) and lower values in developed countries (decelerated aging).

The considered 12 epigenetic characteristics (6 original clocks, 5 PC clocks, DunedinPACE) are not always consistent with each other, so we proposed to calculate an aggregated characteristic describing epigenetic age acceleration as a whole. We applied a voting method in which all epigenetic models “vote” for positive or negative age acceleration, and the “votes” of all epigenetic estimates are equal to each other. We called this metric *Aggregated Sign of <EAA>*, and its value is calculated as the number of epigenetic estimates with positive acceleration minus the number of epigenetic estimates with negative acceleration. We plotted the values of this metric on a map for all available countries (Figure 3D). Aggregated age acceleration varies greatly across continents, countries, and even in neighboring countries, so let’s take a deeper look at the most interesting countries.

Brazil has an Aggregated Sign of <EAA> value of -4 (there are 4 more epigenetic estimates showing decelerated aging than epigenetic estimates showing accelerated aging). For this country, there is only one dataset (45 samples, 50-70 years old) with only women in menopause, in good physical shape (exercising regularly), healthy, non-smokers, not drinking coffee before blood donation, with normal glucose levels [57]. It is worth concluding that a dataset of 45 very healthy women is not representative enough for the entire Brazilian population. Bolivia has an Aggregated Sign of <EAA> value of -2 (there are 2 more epigenetic estimates showing decelerated aging than epigenetic estimates showing accelerated aging). There is only one dataset for this country (37 samples, 37-90 years old) with Tsimane Amerindians. Tsimanes are especially interesting to aging researchers because they experience high levels of inflammation due to recurrent infections, but show minimal risk factors for cardiovascular diseases or type 2 diabetes with age; they have minimal hypertension and obesity, low LDL cholesterol, and no evidence of peripheral arterial disease [33]. It is worth concluding that the dataset of a unique, almost isolated population is not representative enough for the entire Bolivian population. Australia has an Aggregated Sign of <EAA> value of +6 (there are 6 more epigenetic estimates showing accelerated aging than epigenetic estimates showing decelerated aging). There are 2 datasets (640 samples, 4-75 years) for this country. The larger dataset (614) contains twins and their family members for whom the risk of melanoma was tracked (not strict criteria for selecting healthy individuals for the control group, who are still considered healthy in the literature) [58]. The smaller dataset (26) contains 4-year-old children who are in a control group in terms of absence of food allergy [59]. Singapore has an Aggregated Sign of <EAA> value of +4 (there are 4 more epigenetic estimates showing accelerated aging than epigenetic estimates showing decelerated aging). For this country, there is only one dataset (915 samples, 19-46 years) with healthy but early pregnant women only [60]. The inclusion of this dataset in the analysis may be questionable, but we decided to analyze it because it is the only dataset with Singapore residents. Russia has an Aggregated Sign of <EAA> value of -2 (there are 2 more epigenetic estimates showing decelerated aging than epigenetic estimates showing accelerated aging). There is only one dataset (245 samples, 11-101 years) for this country. This dataset has strict exclusion criteria: acute chronic diseases, cancer and respiratory infection at the time of biomaterial donation [48]. Here, the controls are really healthy individuals, so overall epigenetic age acceleration is small in them. Congo has an Aggregated Sign of <EAA> value of +6 (there are 6 more epigenetic estimates showing accelerated aging than epigenetic estimates showing decelerated aging). There are 2 datasets (190 samples, 3-42 years) for this country. The larger dataset (177) contains only women who have recently given birth [61], the smaller dataset contains 3-year-olds [62]. South Korea has an Aggregated Sign of <EAA> value of -6 (there are 6 more epigenetic estimates showing decelerated aging than epigenetic estimates showing accelerated aging). For this country, there is only one dataset (50 samples, 20-56 years old) with strict exclusion criteria: IQ<70, history of head trauma, serious neurological disorders (epilepsy, stroke, Parkinson’s disease and/or dementia) and serious diseases [63].

We also, for each country, divided all samples into females and males, and applied the Mann-Whitney U-test for these two groups, obtaining FDR-adjusted p-values. We then calculated the number of epigenetic clocks that have higher epigenetic age acceleration in males than females (blue bars in Figure 3E), higher epigenetic age acceleration in females than males (red bars in Figure 3E), and no statistical significance/one sex (gray bars in Figure 3E). Males have a higher epigenetic age acceleration than females in most countries. Females have a higher epigenetic age acceleration than males in Australia and Congo. In Singapore, Chile, Brazil and Bolivia, there are either no statistically significant differences between males and females on any epigenetic estimate or the country is represented by only one sex.

The racial/ethnic population composition of different countries can vary greatly. For example, samples from the UK include English, Indians, Scots, and Irish, samples from the USA include African Americans, White Americans, Latinos, Asian Americans, and samples from Singapore include Indians, Chinese, and Malays. Many countries are represented by one population: Chinese in China, Swedes in Sweden, Spanish in Spain, and so on. The relationship between individual datasets, populations, and countries is presented in Figure 4A. A deeper study of epigenetic age acceleration in different populations is also of interest because it provides more detailed insight into racial/ethnic differences in aging rates.

The considered DNA methylation data of blood show even higher diversity of populations compared to countries: 22 countries are represented by 29 populations (Figure 4B). The largest number of samples belong to English (more than 4000 samples), Indians and African Americans (more than 2800 samples in each group). The age range of samples from different populations, as for countries, varies greatly. The widest age ranges are observed for English, African Americans, White Americans, Swedes, Russians, Yakuts; Estonians, Finns, Danes are divided into two subgroups; all samples of Canadians and Flemings are children; Chileans and Koreans are mostly young people 20-40 years; representatives of Dutch are elderly (60-80 years).

Next, we analyzed the mean values of epigenetic age acceleration for different populations and epigenetic models in the form of a diagram with hierarchical clustering (Figure 4C). This diagram may partially be similar to the one for countries (Figure 3B), however, there are some important differences: (1) There are populations that are included in several countries (e.g., Indians are in UK and Singapore, Chinese are in China and Singapore); (2) There are countries that include several populations (e.g., USA is represented by African Americans, White Americans, Latinos, Asian Americans; Russia is represented by Russians, Yakuts). In the first case, an analysis of epigenetic age acceleration would combine the influence from the geographic component and emphasize the influence of genetic similarity between samples, while in the second case, an analysis of epigenetic age acceleration would allow for a more detailed investigation of population differences within countries. As in the case of countries, there is no single population for which all epigenetic clocks show only positive or negative age acceleration. As in the case of countries, for populations PhenoAge and PC-PhenoAge almost always give negative epigenetic age acceleration, while GrimAge2 and PC-GrimAge, on the contrary, give positive age acceleration. When comparing the diagram for countries with the one for populations, several curious differences can be observed. In particular, Singapore, which is in the same cluster as Congo, is represented by three populations. Chinese and Malays remain close to Congolese, while Indians (more numerous in the UK than in Singapore) are in the same cluster as Chileans, Danes and Swedes (where UK is located on the country plot). African Americans, White Americans, Asian Americans (from USA) are close together (about where USA is on the country plot), while Latinos are in the same cluster as Spanish and Italians. Both Russians and Yakuts representing Russia are quite similar to each other, all epigenetic clocks show the same sign of age acceleration for both populations. The most representative countries in terms of population size and population diversity will be investigated in detail in Section 2.4.

We analyzed DunedinPACE separately (mean values for the considered populations and distributions of values are shown in Figure 4D). The slowest aging populations correspond to the slowest aging countries. Most populations have aging rates close to 1. For these populations, the DunedinPACE distribution is quite wide, but the mean value is close to 1. Chinese and Malays age faster than Indians; White Americans age slightly slower than African Americans, Latinos, Asian Americans; Yakuts age faster than Russians. Congolese, Malays and Tsimanes age faster than all other considered populations.

We also compared the aging rate across populations in males and females (Figure 4E). Overall, there are many similarities between countries and populations. Among the interesting differences is that Chinese females age faster than males (in China males age faster, while in Singapore, which also has Chinese, there is no statistically significant difference). In Russians, males age faster than females, while in Yakuts there is no statistically significant difference.

In other tissues, epigenetic age acceleration may also differ between countries and populations. We summarize the main points here. In the analysis of buccal data for representatives of Canada, Netherlands, UK, USA pan-tissue epigenetic models show low values of age acceleration, for representatives of Jamaica all models consistently show high positive age acceleration. Interestingly, in Canada and Netherlands, the majority of samples are children, and girls have higher age acceleration than boys. A notable feature of the brain methylation data is that for all populations, all pan-tissue epigenetic models show negative mean epigenetic age acceleration and a monotonic decrease with age. Monotonically decreasing with age of mean epigenetic age acceleration is also observed in the lung and breast data. The peculiarity of breast methylation data is also that for all populations, all pan-tissue epigenetic models show positive mean epigenetic age acceleration (but since there is only one dataset for this tissue, this may be due to dataset characteristics). In the analysis of colon SkinBloodAge and PC-SkinBloodAge data for all countries and populations demonstrates strongly positive mean epigenetic age acceleration. Detailed results are presented in Supplementary Materials, Supplementary Figures S5-S11.

### 2.4. Epigenetic age acceleration inside the countries

Most countries and populations in the analyzed blood DNA methylation data are represented by a few datasets, but there are large countries for which many samples from multiple datasets are available, and the epigenetic differences between them would be interesting to investigate. We have considered separately three countries for which the largest number of samples are available - UK, USA and Sweden. For them, we have analyzed in detail epigenetic age acceleration in different populations and even datasets. Let us focus on each of them in detail. We started with a detailed analysis of the UK data: during the GEO analysis, 11 datasets were selected with representatives of 4 populations - English, Indians, Scots, and Irish. The numbers of samples in each dataset and in each population with their age distributions are presented in Figure 5A. All datasets have fairly wide age distributions except two, GSE105018 and GSE154566, which contain only 18-year-old participants. The English and Indians populations are the most representative, and all populations have fairly broad age distributions. We calculated the moving average (within 5 years) of epigenetic age acceleration and plotted it as a function of real age (Figure 5B) to investigate how different epigenetic models work at different age ranges and to analyze similarities and differences between populations. Most epigenetic models show similar trends - almost monotonically decreasing epigenetic age acceleration with age (Horvath, GrimAge, GrimAge2, PC-Hannum, PC-Horvath, PC-SkinBloodAge, PC-GrimAge). Different behavior is observed for SkinBloodAge and PhenoAge - for all populations there are “fluctuations” around some age acceleration level (SkinBloodAge in Scots - about 10-12 years, SkinBloodAge in other populations - about 2-5 years, PhenoAge in Scots - about 5 years, PhenoAge in other populations - about -10 years), no significant increases/decreases are observed. Most of the original models show the highest epigenetic age acceleration for Scots compared to other populations (SkinBloodAge, PhenoAge, GrimAge, GrimAge2). An interesting behavior is shown by English for Hannum - the average epigenetic age acceleration first decreases up to 40 years, then increases up to 60-70 years and decreases again at older ages (although it is worth noting that the values themselves do not show very high or low values). DunedinPACE aging rate increases with age in all populations. The mean epigenetic age acceleration for all datasets and populations with hierarchical clustering is shown in Figure 5C. The small dataset which is part of the twin study (GSE154566) has very large predicted values of GrimAge, GrimAge2 and PhenoAge, while PC-modifications and the rest of the clocks show adequate results. This means that GrimAge, GrimAge2 and PhenoAge may be vulnerable to machine learning attacks [64], i.e. a relatively small deviation in the input data drastically degrades the quality of the model, while PC-modifications of the clock are relatively less susceptible to this. Irish have lower mean epigenetic age acceleration for all 12 estimates than the other populations. In English, Indians, and Scots, the mean values of epigenetic acceleration generally show a similar trend: high age acceleration in GrimAge2, PC-GrimAge, PC-Hannum, and SkinBloodAge, negative age acceleration in PC-PhenoAge, and lower absolute values in the other clocks. Separately, we considered the mean DunedinPACE aging rate (Figure 5D) - all datasets and populations show values close to 1. In the largest GSE55763 dataset, DunedinPACE has a maximum (1.059). Irish and English have the minimum DunedinPACE among populations (0.934 and 0.951, respectively), Indians age the fastest (1.095). For most datasets, males have higher epigenetic age acceleration than females (Figure 5E); only GSE105018 (equal number of epigenetic estimates demonstrating accelerated aging of males relative to females) and GSE152027 (females age faster than males) differ. For populations, the pattern is quite homogeneous - all males have higher epigenetic age acceleration than females. The datasets represent residents from different parts of the UK, with a predominance of London residents (Figure 5F).

**Figure 5.**
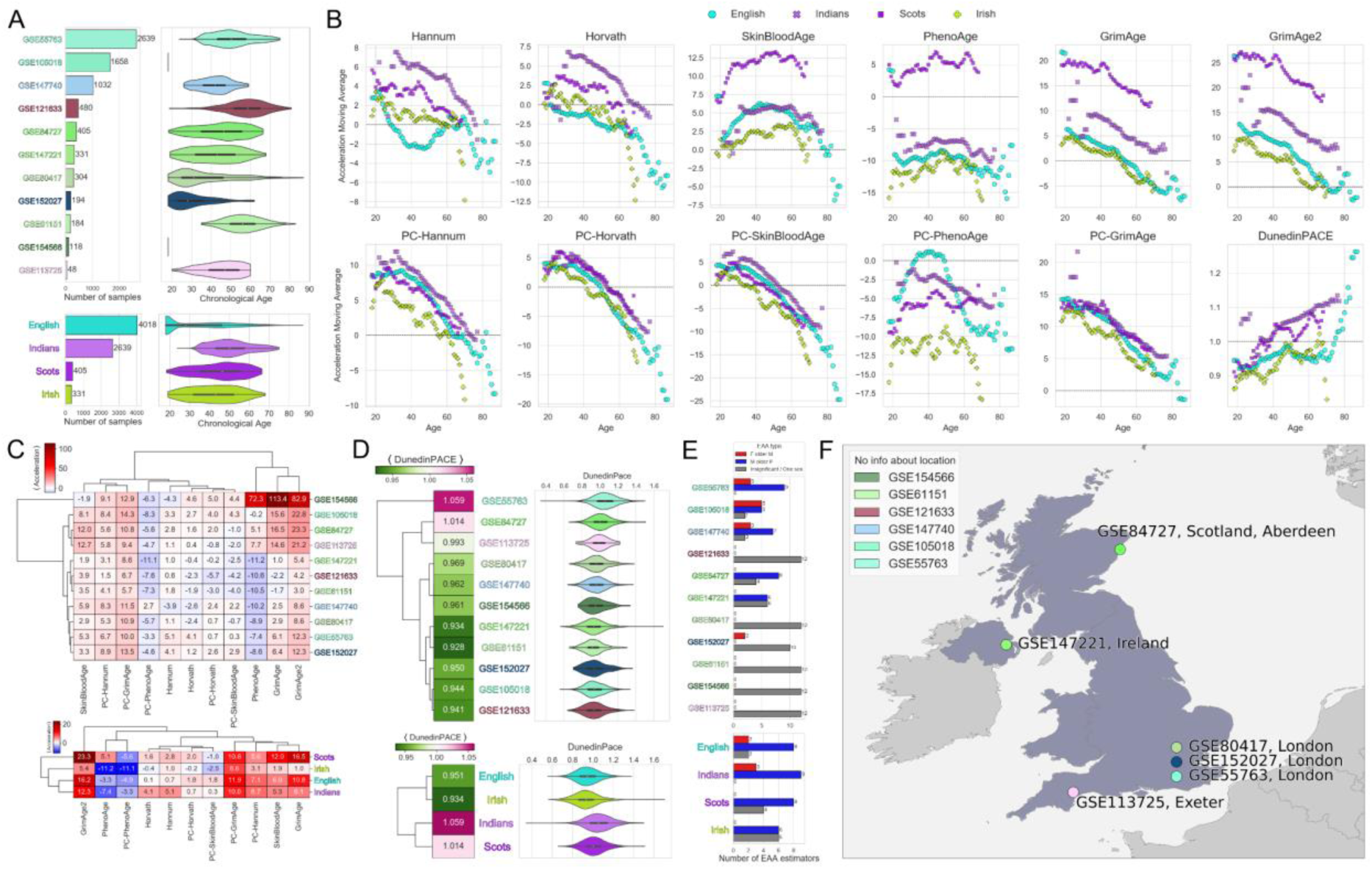
Epigenetic age acceleration in different datasets (GSEs) and populations in the UK. (A) Number of samples from different datasets (top left) and populations (bottom left) with corresponding age distributions (top right and bottom right, respectively). (B) Dependence of the moving average (within 5 years) of epigenetic age acceleration on real age for all populations. (C) Diagram of the mean values of epigenetic age acceleration for datasets (top) and populations (bottom) with hierarchical clustering. (D) Diagram of the mean values of DunedinPACE for datasets (top) and populations (bottom) with hierarchical clustering. (E) Number of epigenetic metrics for each dataset (top) and populations (bottom) for which: males have higher epigenetic age acceleration (blue), females have higher epigenetic age acceleration (red), no statistical significance/one sex (gray). Mann-Whitney U-test was applied for males and females for each tissue, with FDR-corrected resulting p-values. (F) Map of the UK with marked locations of samples from different datasets. The upper left corner indicates datasets for which the exact location of samples within the UK is unknown.

We next performed a detailed analysis of the USA: during the GEO analysis, we selected 20 datasets (the largest number among all countries) that contain representatives of 4 populations - African Americans, White Americans, Latinos, and Asian Americans. The numbers of samples in each dataset and in each population with their age distributions are presented in Figure 6A. Most of the datasets have fairly broad age distributions, GSE132181 (only 7-year-olds), GSE219037 and GSE227809 (about 20 years old), and GSE107459 differ. African Americans and White Americans are the most representative populations; all populations except Asian Americans have rather broad age distributions. We calculated a moving average (within 5 years) of epigenetic age acceleration and plotted it as a function of real age (Figure 6B) to investigate how epigenetic models work at different age ranges and to analyze similarities and differences between populations. As for the UK, many epigenetic models show an almost monotonic decrease in epigenetic age acceleration with age (Horvath, GrimAge, GrimAge2, PC-Hannum, PC-Horvath, PC-SkinBloodAge, PC-GrimAge). According to Hannum, different populations show different behavior - African Americans and Latinos show a decrease in epigenetic age acceleration with age, while White Americans, on the contrary, show an increase until the age of 50, after which epigenetic acceleration values fluctuate around 0. Different behavior is also observed for SkinBloodAge and PhenoAge - for all populations there are “fluctuations” around some level (SkinBloodAge around 0-5 years, PhenoAge around -10 years), no significant increases/decreases are observed. Overall, the entire USA cohort appears to be fairly homogeneous, with no population that stands out significantly on any of the epigenetic estimates. DunedinPACE aging rate increases with age in all populations, only the starting point differs, which is highest for African Americans and lowest for White Americans. Mean epigenetic age acceleration for all datasets and populations with hierarchical clustering is presented in Figure 6C. Systematically low acceleration values are shown by PhenoAge and PC-PhenoAge, isolated in a separate hierarchical cluster. GrimAge2 and PC-GrimAge have maximum values. Other epigenetic ages have no clear tendencies to overestimate/underestimate acceleration. For populations, the mean values of epigenetic acceleration generally have a similar trend: high age acceleration for GrimAge2, PC-GrimAge, PC-Hannum, SkinBloodAge, negative acceleration for PhenoAge, PC-PhenoAge, PC-Horvath, PC-SkinBloodAge (separate group in hierarchical clustering). The remaining clocks have smaller absolute values. Overall, there is relative consistency between datasets and populations for all epigenetic estimates of accelerated aging in the US, in contrast to the UK. We considered the DunedinPACE mean aging rate separately (Figure 6D). The datasets vary quite widely in aging rate values. The systematically low values of age acceleration and DunedinPACE in the GSE147430 dataset may be due to the choice of control group (they are non-smokers, have clean medical records and no acute diseases) [65]. In contrast, datasets GSE174818 (control group has a negative test for COVID-19) [66], GSE72680 (control group is not taking any treatment but may have childhood trauma) [67] and GSE77696 (control group are veterans without HIV) [68] have higher age acceleration and DunedinPACE values. The spread of mean DunedinPACE values is quite large, ranging from 0.62 for GSE147430 to 1.23 for GSE174818. Relative to populations, DunedinPACE is minimal in White Americans and maximal in African Americans, but generally close to 1 in all populations. For all datasets, either males have more epigenetic age acceleration than females or there is no statistical significance/one sex (Figure 6E). For populations, the pattern is homogeneous, with all males having higher epigenetic age acceleration than females. Datasets represent residents from the western, southern, and eastern parts of the USA; residents from the northern and central parts are not represented (Figure 6F).

**Figure 6.**
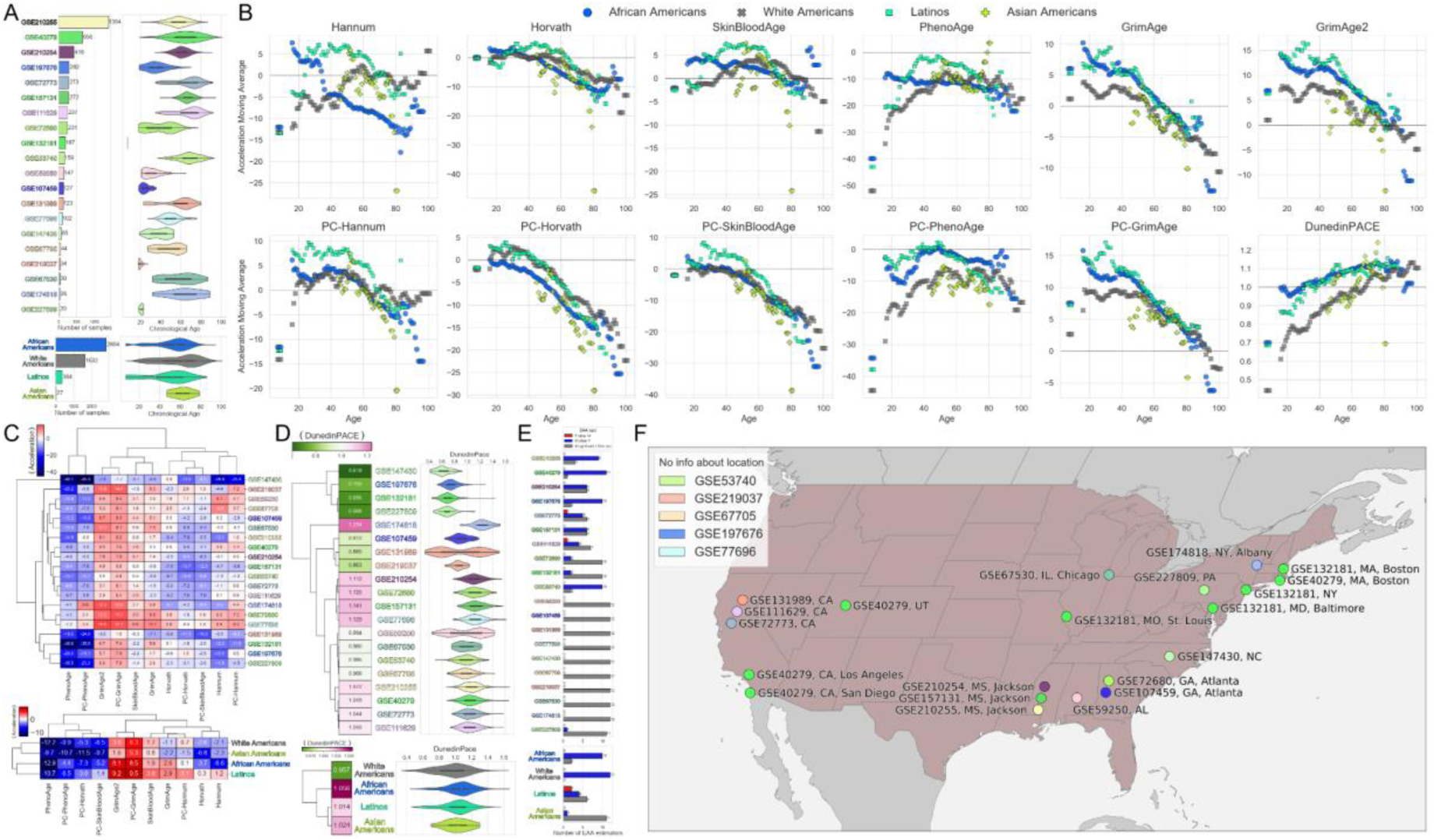
Epigenetic age acceleration in different datasets (GSEs) and populations in the USA. (A) Number of samples from different datasets (top left) and populations (bottom left) with corresponding age distributions (top right and bottom right, respectively). (B) Dependence of the moving average (within 5 years) of epigenetic age acceleration on real age for all populations. (C) Diagram of the mean values of epigenetic age acceleration for datasets (top) and populations (bottom) with hierarchical clustering. (D) Diagram of the mean values of DunedinPACE for datasets (top) and populations (bottom) with hierarchical clustering. (E) Number of epigenetic metrics for each dataset (top) and populations (bottom) for which: males have higher epigenetic age acceleration (blue), females have higher epigenetic age acceleration (red), no statistical significance/one sex (gray). Mann-Whitney U-test was applied for males and females for each tissue, with FDR-corrected resulting p-values. (F) Map of USA with marked locations of samples from different datasets. The upper left corner shows datasets for which the exact location of samples within the USA is unknown.

Another country that we analyzed in detail is Sweden: 4 datasets were selected during the GEO analysis, all representing the same population, Swedes. The numbers of samples in each dataset with their age distributions are shown in Figure 7B. The 3 datasets have wide age distributions, while the dataset GSE73103 has only young samples (15-35 years old). We remind that the GSE87571 dataset is the reference dataset for harmonization of blood DNA methylation data [69]. We calculated a moving average (within 5 years) of epigenetic age acceleration values and plotted it as a function of real age (Figure 7A) to investigate how epigenetic models perform at different age ranges. All datasets in Sweden are homogeneous and behave similarly across all epigenetic estimates. Most epigenetic models show almost monotonic decreasing epigenetic age acceleration with age (Hannum, Horvath, GrimAge, GrimAge2, PC-Hannum, PC-Horvath, PC-SkinBloodAge, PC-GrimAge). At the same time in Hannum, PC-Hannum, PC-GrimAge, GrimAge2 epigenetic age acceleration is positive almost on the whole age range and becomes negative only in old age (more than 80 years), in other epigenetic models with monotonous behavior about 60 years there is a transition from positive to negative zone. The behavior on SkinBloodAge, PhenoAge and PC-PhenoAge differs - for all datasets, there is first an increase, then a decrease. DunedinPACE aging rate increases with age, crossing 1 around 60 years. The mean epigenetic age acceleration for all datasets and populations with hierarchical clustering is presented in Figure 7C. As in the previous cases, PhenoAge with its PC-version show a rather strong negative age acceleration, while GrimAge2 and PC-GrimAge have a large positive age acceleration. PC-Hannum is also slightly higher compared to the original Hannum clock model. All datasets are quite homogeneous in terms of age acceleration values, the metrics in all datasets have the same sign. We separately considered the mean DunedinPACE aging rate (Figure 7D), which varies significantly between datasets with GSE73103 having a mean value close to 0.8 (may be due to the fact that DunedinPACE often underestimates for young samples), while GSE87571 and GSE42861 are around 0.97. In general, DunedinPACE values in Sweden are less than 1 (corresponding to decelerated aging). The datasets represent residents from northern and eastern Sweden (Figure 7E).

**Figure 7.**
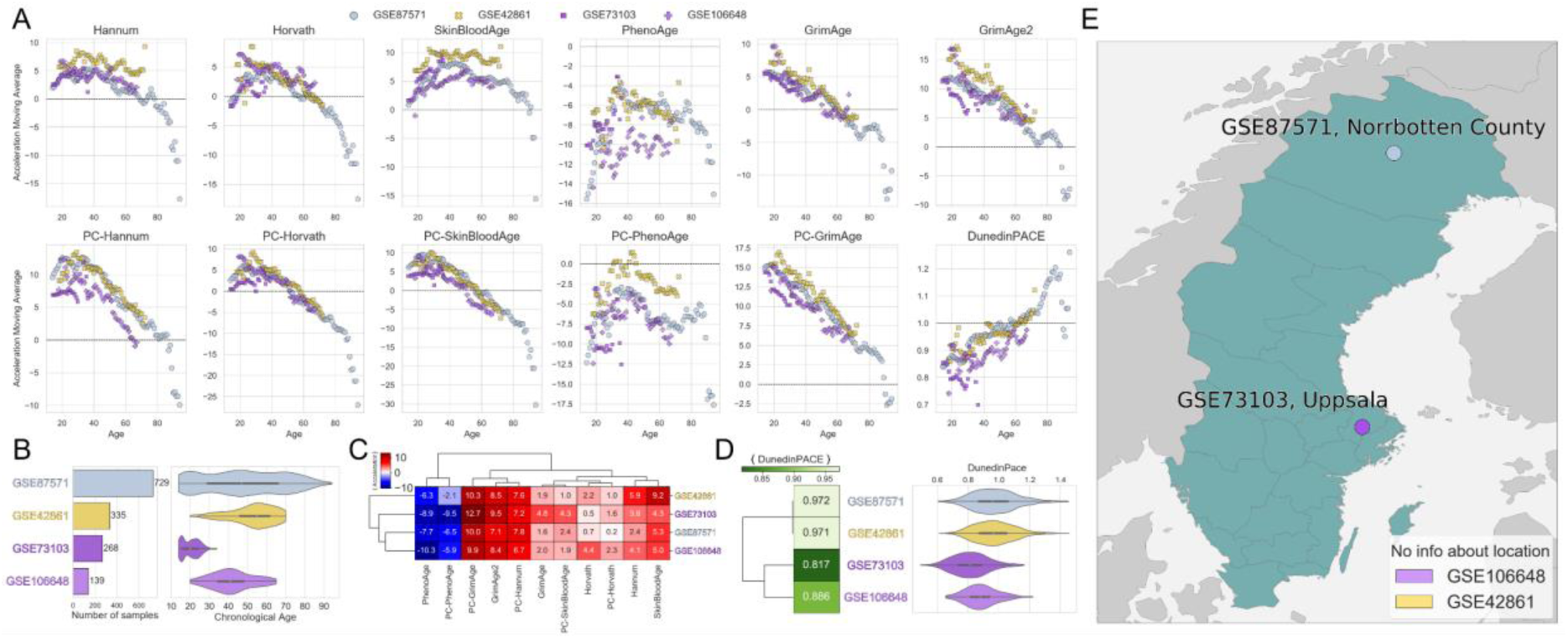
Epigenetic age acceleration in different datasets (GSEs) in Sweden. (A) Dependence of the moving average (within 5 years) of epigenetic age acceleration on real age for all datasets. (B) Number of samples from different datasets (left) with corresponding age distributions (right). (C) Diagram of the mean values of epigenetic age acceleration for datasets with hierarchical clustering. (D) Diagram of the mean values of DunedinPACE for datasets with hierarchical clustering. (E) Map of Sweden with marked locations of samples from different datasets. The bottom right corner shows datasets for which the exact location of samples within Sweden is unknown.

## 3. Discussion

### 3.1. Conclusion

In this work, we performed a large study of epigenetic age acceleration in different parts of the globe, the meta-analysis included open access DNA methylation data from the GEO repository, taken from 10 tissue groups, representing 25 countries and 31 populations, with a separate focus on the sex-specific patterns of age acceleration. We investigated in detail the most popular models of epigenetic clock and aging rates, and compared their behavior across tissues, countries, races/populations, sexes, and age ranges. It is important that only healthy controls are analyzed to avoid the influence of different diseases on epigenetic age acceleration.

The most common tissue that has DNA methylation data is blood, more than 10 times fewer data are available for buccal and brain, all other tissues are even fewer. Although most epigenetic estimates models are trained on blood DNA methylation data, we were interested in highlighting the differences in DNA methylation of different tissues and thus in epigenetic age acceleration for them. Most epigenetic clocks (Hannum, Horvath, GrimAge, all PC clocks, GrimAge2) for almost all tissues show a decrease in mean epigenetic age acceleration with age. Only SkinBloodAge and PhenoAge differ, which for some tissues (brain, muscles) also show a decrease in average epigenetic age acceleration with age, while for most tissues there is no stable dependence on age (either fluctuations around a certain level or non-monotonic changes with age are observed). It is natural that all epigenetic models work best for blood, and the pan-tissue Horvath epigenetic clock model handles almost all tissues. All versions of GrimAge (GrimAge, GrimAge2, PC-GrimAge) for all investigated tissues demonstrate positive mean age acceleration, while PhenoAge for most tissues (except saliva, buccal and colon) demonstrates negative mean age acceleration. According to DunedinPACE (constructed for blood methylation), only blood and has an aging rate of about 1 (one biological year per one chronological year), all other tissues age accelerated. There is no tissue for which the number of epigenetic clocks for which females age faster than males is higher than the number of epigenetic clocks for which males age faster than females.

The detailed analysis of the representation of blood DNA methylation data by country has shown that it is quite broad - the analysis includes representatives of countries on all continents. The age range of data from different countries varies considerably: somewhere it is very wide and represents most age groups from the youngest to the oldest (e.g., in the USA, Sweden, Russia), and somewhere - on the contrary, it is very narrow and contains only children or only the elderly. All epigenetic models work differently for different countries, and there is no country for which all epigenetic models consistently show only positive or only negative mean epigenetic age acceleration. GrimAge2 and PC-GrimAge systematically show positive age acceleration for almost all countries, while PhenoAge and PC-PhenoAge, on the contrary, show negative age acceleration. The DunedinPACE aging rate demonstrates reasonable and adequate results: very low values of the average aging rate for children (Canada, Belgium), normal aging rate (about 1) for most developed countries with a sufficiently high standard of living (most European countries, USA), higher aging rate values for the rest of the countries (Russia, Bolivia, Congo). For a more aggregated representation of the performance of different epigenetic models in different countries, we introduced a special metric, Aggregated Sign of EAA, which is based on the “voting” of all metrics for “positive” or “negative” age acceleration. Such a metric allows to reveal very interesting patterns - for example, to verify the dependence of epigenetic age acceleration on the characteristics of a particular dataset, which may not always fully characterize the population of the whole country. In particular, low epigenetic age acceleration in Brazil or high epigenetic age acceleration in Australia is due to the characteristics of the datasets (Brazil has only healthy, regularly exercising women, Australia has a general population for which melanoma risk is being studied). This metric makes it clear that the problem of different criteria for defining a control group (healthy people without acute and chronic disease, people without the specific disease being studied, the general population without severe disease, and so on) is acute here. In most countries, more epigenetic models show higher epigenetic age acceleration in males, but in Australia and Congo more models show higher epigenetic age acceleration in females.

Although many countries are represented by only one racial/ethnic population, there are examples of countries with members of more than one population, as well as examples of populations living in more than one country. As with countries, for different populations, epigenetic models show inconsistent results - there are no populations for which all epigenetic models show age acceleration of the same sign. GrimAge2 and PC-GrimAge, for almost all populations, systematically show positive age acceleration, while PhenoAge and PC-PhenoAge, on the contrary, show negative age acceleration. African Americans, White Americans and Asian Americans living in the USA are similar in terms of mean epigenetic age acceleration according to different estimates, while Latinos are closer to Italians and Spanish. DunedinPACE aging rate also differs between populations living in the same country - for example, Russians living in Russia have a lower aging rate than Yakuts living in the same country, but in more severe climatic conditions. If we look at populations, women have more epigenetic estimators showing age acceleration not only in White Australians and Congolese but also in Chinese.

We also analyzed epigenetic age acceleration in the UK, USA, Sweden separately for individual populations (if available) and datasets. For all original clocks except Hannum and Horvath, Scots are aging faster than other populations. English over 80 years shows negative age acceleration in all epigenetic clocks. DunedinPACE aging rate is close to 1 in all UK populations. The US populations are more homogeneous and show similar behavior across all epigenetic metrics with age in terms of age acceleration. The pattern is similar for Swedes, but DunedinPACE aging rate is lower than 1. In all countries, males have higher rates of epigenetic age acceleration.

In general, we can conclude that different epigenetic clocks do not agree with each other in the sign of age acceleration for different countries and populations. The difference between the mean values of age acceleration in some pairs of epigenetic clocks can exceed 25-30 years. At the same time, some epigenetic clocks have strong bias: different versions of GrimAge tend to overestimate chronological age, and PhenoAge underestimates it. At the same time, DunedinPACE is a rather reliable characteristic that corresponds well with the standard of living in different countries and shows higher values in developing countries and lower values in developed countries, although it shows extremely low values for children, which is not surprising since the training data included only adults.

Not only geographic and ethnic characteristics affect age acceleration. As shown in Section 2.3, it is important how healthy controls are defined in the data, and epigenetic age acceleration depends strongly on specific datasets in different countries. Where the health criteria are stricter, relatively less age acceleration is observed. However, it is not possible to separate completely the contributions of ’geography/environment’, ’ethnicity’ and ’health’. It is possible to pick out relatively more and relatively less healthy individuals in any country, compared to the training data of the clocks. Small datasets are unlikely to characterize the population as a whole. Ideally, it would be interesting to compare healthy controls selected using the same criteria worldwide, with a broad uniform distribution across countries, but such data are very expensive to collect and cannot be publicly available. Nevertheless, the largest open-access repository of DNA methylation data, GEO, allows us to analyze the mechanisms of epigenetic aging with a focus on geography and ethnicity.

### 3.2. Limitations

The resulting map of epigenetic age acceleration does not always reliably reflect trends of population aging in different countries. Datasets from some countries are not representative. We considered the largest datasets in the GEO repository, but not all, and some unique populations with few samples may still be left out.

The results of the meta-analysis depend heavily on how healthy controls are defined in each individual dataset. In some countries and datasets, the criteria are much stricter than in others (absence of chronic diseases, oncology, diseases and ailments in the acute stage is required), while in other datasets only information on the absence of any disease may be available, with no indication of comorbidities.

Existing epigenetic clocks have some weaknesses - they are very sensitive to data, batch effects, have significant acceleration and deceleration even on conditionally control subjects due to the fact that the data on which they are tested are often significantly different from the training data. As in many areas of machine learning, a rule-of-thumb works: the bigger, more complete, and more diverse the training data, the better-quality metrics the models demonstrate (as, for example, pan-tissue Horvath clock trained on multiple datasets performs relatively well on more than just blood).

The amount of methylation data increases over the years and it is reasonable to update the clock models by expanding their training data so that they perform reliably in a larger number of cases. It also makes sense to use state-of-the-art machine learning models when developing new clocks. ElasticNet, on which most clocks are based, is a good and proven tool for reducing the input feature space. But it is still a linear model, assuming linear relationships between the features. At the same time, modern neural network architectures, such as generalizations of attention and transformer mechanisms for tabular methylation data, are nonlinear and in many studies perform better in terms of model quality metrics [70], and current trends of explainable artificial intelligence will allow a reasonable biological interpretation [71].

## 4. Methods

### 4.1. Epigenetic data processing

We used the R packages GEOmetadb [72], GEOquery [73] to do preliminary parsing of all available datasets in GEO. These tools allow access to metadata associated with samples, platforms, and datasets. The GEOparse python package [74] was used to retrieve characteristics and fields in datasets, and in some cases for automated downloading of preprocessed methylation data.

Next, we describe the details of the selection procedure of datasets.

First, we selected only healthy controls. Since our aim was to compare age acceleration across countries and populations, we needed to exclude the influence of diseases - they are very heterogeneous and can have a significant impact on the final results. Here we should make an important note: healthy controls can be defined differently in each dataset and do not necessarily include absolutely healthy people: in some datasets controls are healthy people without chronic diseases and acute conditions, in some datasets controls are people without certain studied diseases, in some datasets controls are people without bad habits, and so on.

Second, all samples must have information on sex and age. This is necessary to calculate epigenetic age estimates (including using the Horvath’s calculator), as well as epigenetic age acceleration values. Sex is also necessary to perform sex-specific analysis.

Third, samples should have an exact race/ethnicity description so that the meta-analysis would have a complete picture of the compared populations. Samples with unknown race/ethnicity were excluded, and samples with an unknown country of origin/residence were also excluded. Fourth, the age of the participants should be greater than or equal to 3 years. We still decided not to consider newborns and children younger than 3 years of age because the most active development is happening in the first 3 years, after which a new stage is coming [75–78].

Fifth, the dataset should initially contain at least 100 individuals (there may be fewer healthy controls who fit all criteria). We did not consider small datasets because few samples per group may not yield robust statistical results. Because manual preprocessing was performed for all datasets (methylation data may be presented in different formats; datasets may lack some important features on GEO, but these features may be presented in the supplementary materials of related articles), the largest datasets were considered first. One of the options could have been to consider only datasets that have complete information on GEO, but we did not want to lose potentially valuable information, unique population data, so we took a complex but detail-accurate path.

Sixth, the dataset should be released on GEO no later than December 14, 2023.

Additional criteria for excluding samples: (1) Each tissue must have at least 10 samples in total, tissues with fewer samples are not considered; (2) Each ethnic group must have at least 10 samples in total (for all tissue types) - groups with fewer samples are not considered; (3) Each country (state) must have at least 10 samples in total (for all tissue types) - countries with fewer samples are not considered. This exclusion is necessary to ensure that only representative size groups with statistically significant results remain.

### 4.2. Epigenetic ages estimators and statistical analysis

The main details of the used epigenetic models are summarized in Table 1. The results of the epigenetic clocks Hannum DNAmAge [16], Horvath DNAmAge [14], SkinBlood DNAmAge [15], DNAm PhenoAge [17], GrimAge [18], GrimAge version 2 [42] were obtained using Horvath’s DNA Methylation Age Calculator [79]. PC-clocks [41] and DunedinPACE [19] results were obtained using the programming code accompanying the original articles.

Horvath’s DNA Methylation Age Calculator requires specifying a tissue that is selected from a special list and does not always support all detailed tissue types. Therefore, in the meta-analysis we considered aggregated tissue groups - all blood-related tissue types (whole blood, peripheral blood, specific cell types, etc.) were combined into one group “Blood”, all brain-related tissue types (different parts of the brain) were combined into one group “Brain”, and so on.

To analyze the mean epigenetic age acceleration in different age groups, we calculated the moving average in a window of 5 years.

Among the considered 12 epigenetic characteristics (6 original clocks + 5 PC clocks + 1 DunedinPACE) there is no the most important or, on the contrary, the most unimportant clocks. But they do not always agree with each other and it would be clearer to get an aggregated picture describing the epigenetic age acceleration as a whole. Adding up all accelerations of the clocks is not the best option, because there are clocks with large systematic positive (GrimAge2) or negative (PhenoAge, PC-PhenoAge) age acceleration, and their impact will be dominant when averaging over all clocks (it is still unclear how to take into account DunedinPACE, which is not measured in years). For the above reasons, we applied a voting method in which the “votes’’ of all epigenetic estimates are equal to each other. Each epigenetic estimate can show either a positive age acceleration or a negative age deceleration (for epigenetic clock models, the sign of the epigenetic acceleration is taken, and for DunedinPACE, a value less than 1 is considered as a negative deceleration, a value greater than 1 is considered as a positive acceleration). Next, we can calculate the number of “positively” and “negatively” accelerated clocks and introduce the *Aggregated Sign of <EAA= = Num positive - Num negative* (Figure 3D). This characteristic takes values from -12 (all epigenetic ages and DunedinPACE show negative age acceleration) to +12 (positive age acceleration in all clocks and DunedinPACE).

Female and male groups and pairwise comparisons of tissues and countries (Supplementary Figures S1-S4) were performed using the Mann-Whitney U-test [56]. All p-values were FDR-corrected using the Benjamini-Hochberg procedure [80].

## Data availability statement

No new data was generated. Data used in this study are available from the GEO database under accession numbers: GSE55763 [81], GSE105018 [82], GSE84727 [83], GSE51032, GSE152027 [84], GSE87571 [69], GSE147221 [84], GSE125105 [85], GSE42861 [86], GSE74193 [87], GSE80417 [83], GSE40279 [16], GSE56105 [58], GSE111629 [88], GSE121633 [89], GSE59250 [90], GSE72680 [67], GSE210254 [91], GSE53740 [92], GSE112893 [93], GSE73103 [94], GSE90124 [95], GSE61496 [96], GSE72773 [33], GSE59065 [97], GSE67705 [98], GSE106648 [99], GSE157131 [100], GSE125895 [101], GSE137898 [102], GSE111223 [88], GSE137903 [102], GSE137688 [102], GSE110128 [103], GSE80261 [104], GSE124366 [105], GSE113725 [106], GSE61151 [107], GSE69270 [108], GSE74548 [109], GSE75704 [110], GSE74414 [111], GSE116379 [112], GSE41826 [113], GSE67530 [114], GSE89707 [115], GSE61256 [116], GSE71955 [117], GSE107459 [118], GSE68194 [119], GSE101961 [120], GSE147430 [65], GSE94876 [121], GSE85566 [122], GSE201872 [123], GSE103027 [124], GSE77696 [68], GSE131989 [125], GSE115278 [126], GSE210255 [91], GSE147740 [127], GSE158063 [60], GSE179325 [128], GSE219037 [129], GSE175458 [130], GSE132181 [131], GSE220622 [132], GSE185061 [133], GSE224363 [61], GSE197676 [134], GSE234461 [48], GSE207927 [135], GSE147040 [136], GSE172365 [137], GSE171140 [138], GSE227809 [139], GSE224573 [62], GSE143157 [140], GSE164822 [141], GSE137682 [102], GSE118144 [142], GSE164056 [143], GSE216024 [144], GSE157252 [63], GSE174818 [66], GSE196432 [145], GSE201752 [146], GSE151485 [147], GSE199700 [57], GSE154566 [148], GSE151732 [149], GSE114134 [59], GSE142257 [150].

## Code availability statement

No new or unpublished methods were used in the study.

## Author’s Contributions

Conceptualization: I.Y., A.K.; Methodology: I.Y., A.K.; Formal analysis: I.Y., A.K.; Writing – original draft: I.Y., A.K.; Writing – review & editing: I.Y., A.K., C.F., M.I.; Visualization: I.Y., A.K.; Supervision: C.F., M.I.

## Abbreviations

DNA: DeoxyriboNucleic Acid
DNAm: DNA methylation
EAA: Epigenetic Age Acceleration
EEAA: Extrinsic Epigenetic Age Acceleration
FDR: False Discovery Rate
GEO: Gene Expression Omnibus
GeoEPR: Geo-referencing Ethnic Power Relations
IEAA: Intrinsic Epigenetic Age Acceleration
PC: Principal Component
PCA: Principal Component Analysis
regRCPqn: regional Regression on Correlated Probes with quantile normalization.

