## Supplementary Materials for "Map of epigenetic age acceleration: a worldwide meta-analysis"

### Map of epigenetic age acceleration: a worldwide meta-analysis Supplementary Materials

#### 1. Tissues: statistical pairwise comparison

We made a pairwise comparison of all considered tissues for all epigenetic metrics. For this, we applied the Mann-Whitney U-test for each pair of tissues, the resulting FDR-corrected p-values are shown in the lower triangles of the diagrams in Supplementary Figures S1, S2. The absence of a statistically significant difference is shown in black. In the upper triangles, we plotted the difference in mean epigenetic age acceleration values (or mean DunedinPACE values) for all tissue pairs (in Supplementary Figures S1, S2, the difference is presented as column minus row).

The very small number of black cells indicates that the considered tissues are statistically significantly different in terms of mean epigenetic age acceleration for almost all epigenetic estimates. The absence of a statistically significant difference is most often observed in tissue pairs involving Liver, which may be due to the small sample size for this tissue. The lowest p-values are most often observed for Blood-Buccal, Blood-Brain, and Brain-Buccal pairs.

For PhenoAge and DunedinPACE, all tissue pairs are statistically significantly different. Epigenetic clock models and their PC-modifications do not always give consistent results (e.g., mean differences behave differently for PhenoAge and PC-PhenoAge). The smallest absolute values of the mean epigenetic age acceleration difference are observed for Horvath DNAmAge and its PC-modification.

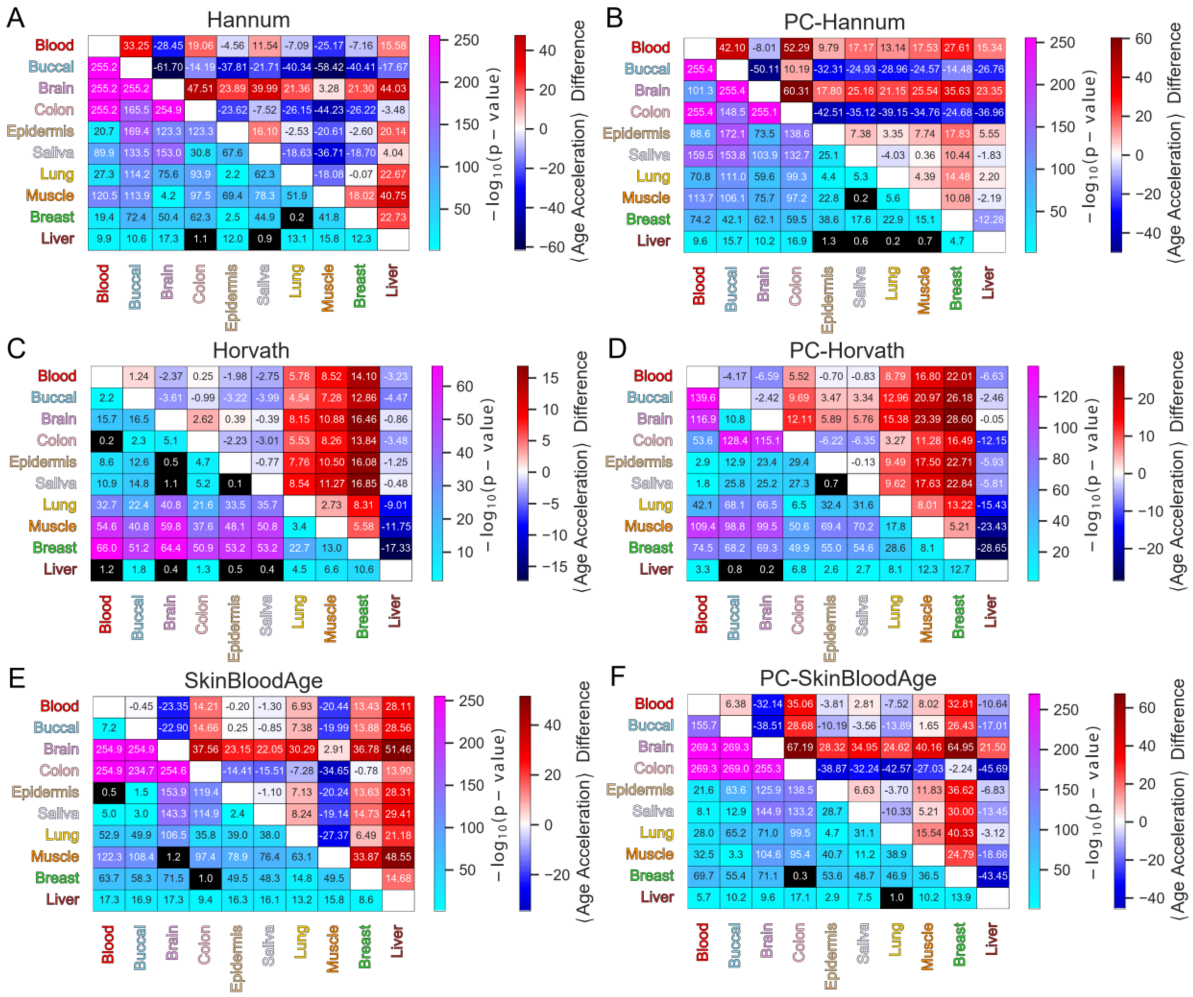


Supplementary Figure S1. Statistical significance of the difference in mean (A) Hannum DNAmAge, (B) PC-Hannum DNAmAge, (C) Horvath DNAmAge, (D) PC-Horvath DNAmAge, (E) SkinBlood DNAmAge, (F) PC-SkinBlood DNAmAge values between different tissues. Lower triangle: FDR-corrected pairwise Mann-Whitney U test p-value. Upper triangle: Difference between mean (A) Hannum DNAmAge, (B) PC-Hannum DNAmAge, (C) Horvath DNAmAge, (D) PC-Horvath DNAmAge, (E) SkinBlood DNAmAge, (F) PC-SkinBlood DNAmAge values (column minus row).

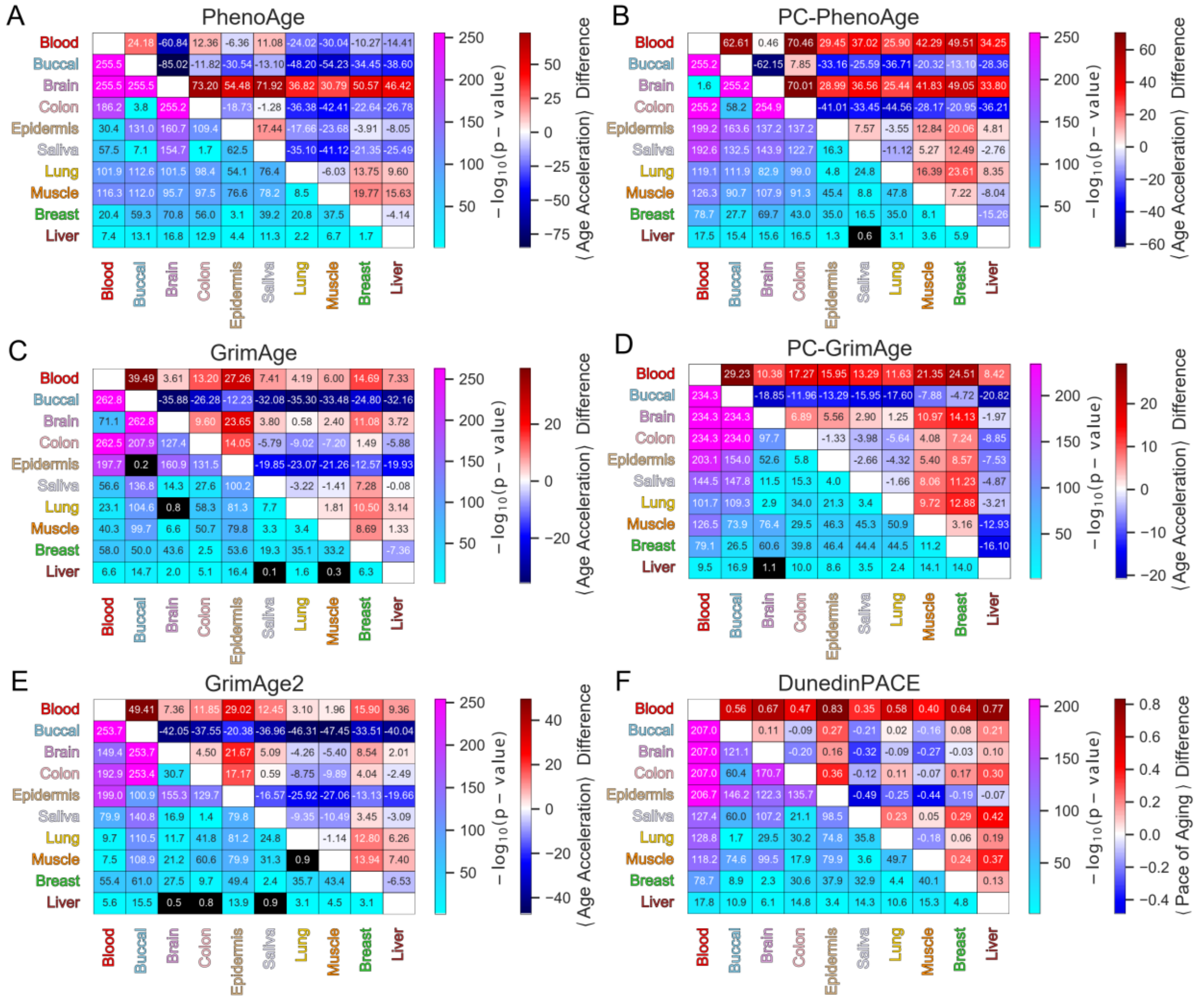


Supplementary Figure S2. Statistical significance of the difference in mean (A) DNAm PhenoAge, (B) PC-PhenoAge, (C) GrimAge, (D) PC-GrimAge, (E) GrimAge2, (F) DunedinPACE values between different tissues. Lower triangle: FDR-corrected pairwise Mann-Whitney U test p-value. Upper triangle: Difference between mean (A) DNAm PhenoAge, (B) PC-PhenoAge, (C) GrimAge, (D) PC-GrimAge, (E) GrimAge2, (F) DunedinPACE values (column minus row).

#### 2. Countries: statistical pairwise comparison for blood

We made a pairwise comparison of all countries with available blood DNA methylation data for all epigenetic metrics. For this, we applied the Mann-Whitney U-test for each pair of countries, the resulting FDR-corrected p-values are shown in the lower triangles of the diagrams in Supplementary Figures S3, S4. The absence of a statistically significant difference is shown in black. In the upper triangles, we plotted the difference in mean epigenetic age acceleration values (or mean DunedinPACE values) for all country pairs (in Supplementary Figures S3, S4, the difference is presented as column minus row).

The rather small number of black cells indicates that the considered countries are mostly statistically significantly different in terms of mean epigenetic age acceleration for almost all epigenetic estimates. The absence of statistically significant difference is most often observed in country pairs involving Chile, Belgium, China, and Brazil, which may be due to the small sample size for these countries (only one or two datasets each). There is no single epigenetic estimate for which all country pairs are statistically significantly different. The smallest p-values are most often observed for the US-UK, US-Sweden, and US-Singapore pairs.

Epigenetic clock models and their PC-modifications do not always give consistent results (e.g., mean differences behave differently for Hannum and PC-Hannum, GrimAge and PC-GrimAge). The smallest absolute values of the mean epigenetic age acceleration difference are observed for the two Horvath's models (Horvath DNAmAge and SkinBlood DNAmAge) and their PC-modifications.

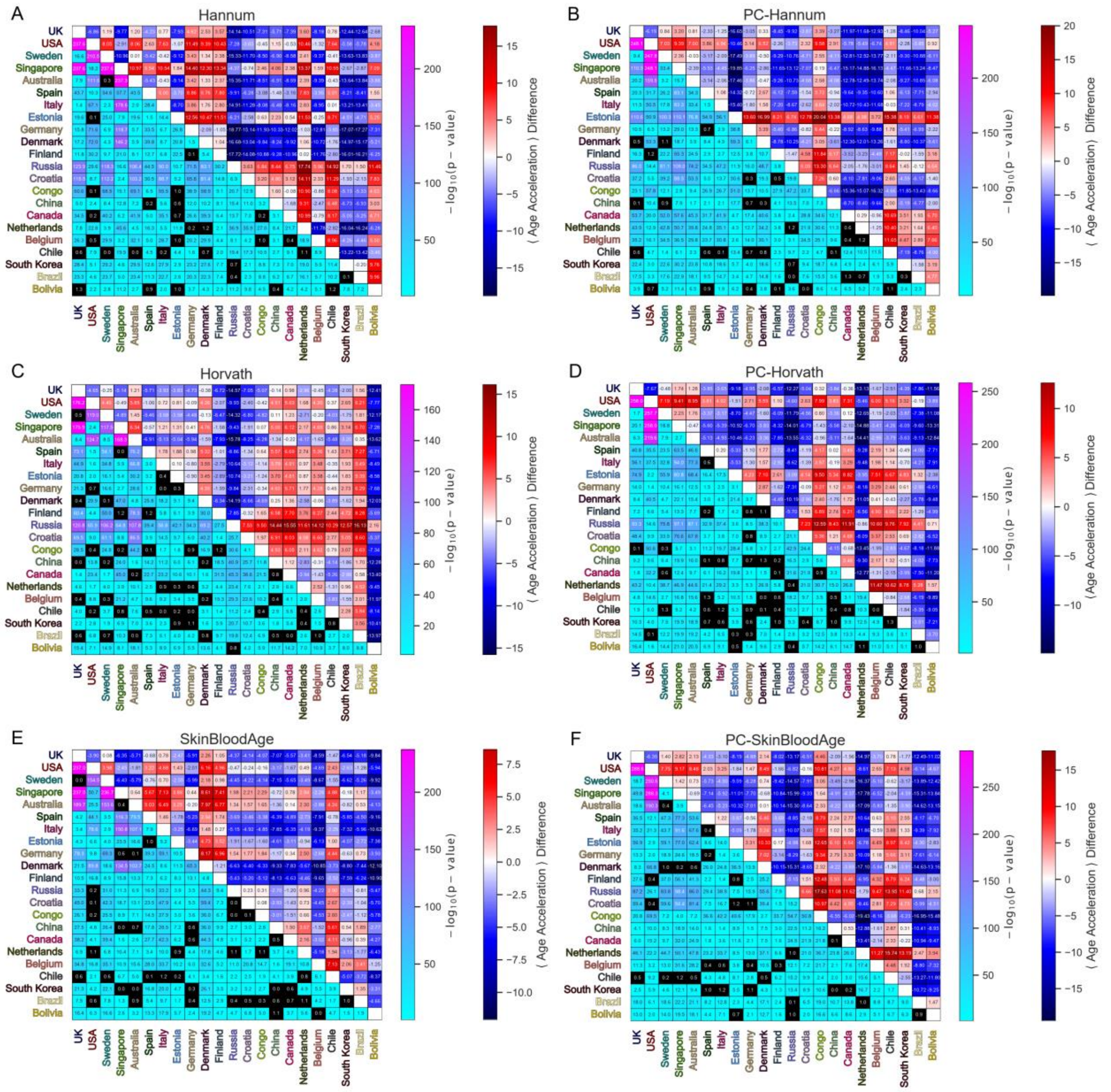


Supplementary Figure S3. Statistical significance of the difference in mean (A) Hannum DNAmAge, (B) PC-Hannum DNAmAge, (C) Horvath DNAmAge, (D) PC-Horvath DNAmAge, (E) SkinBlood DNAmAge, (F) PC-SkinBlood DNAmAge values between different countries for blood. Lower triangle: FDR-corrected pairwise Mann-Whitney U test p-value. Upper triangle: Difference between mean (A) Hannum DNAmAge, (B) PC-Hannum DNAmAge, (C) Horvath DNAmAge, (D) PC-Horvath DNAmAge, (E) SkinBlood DNAmAge (F) PC-SkinBlood DNAmAge values (column minus row).

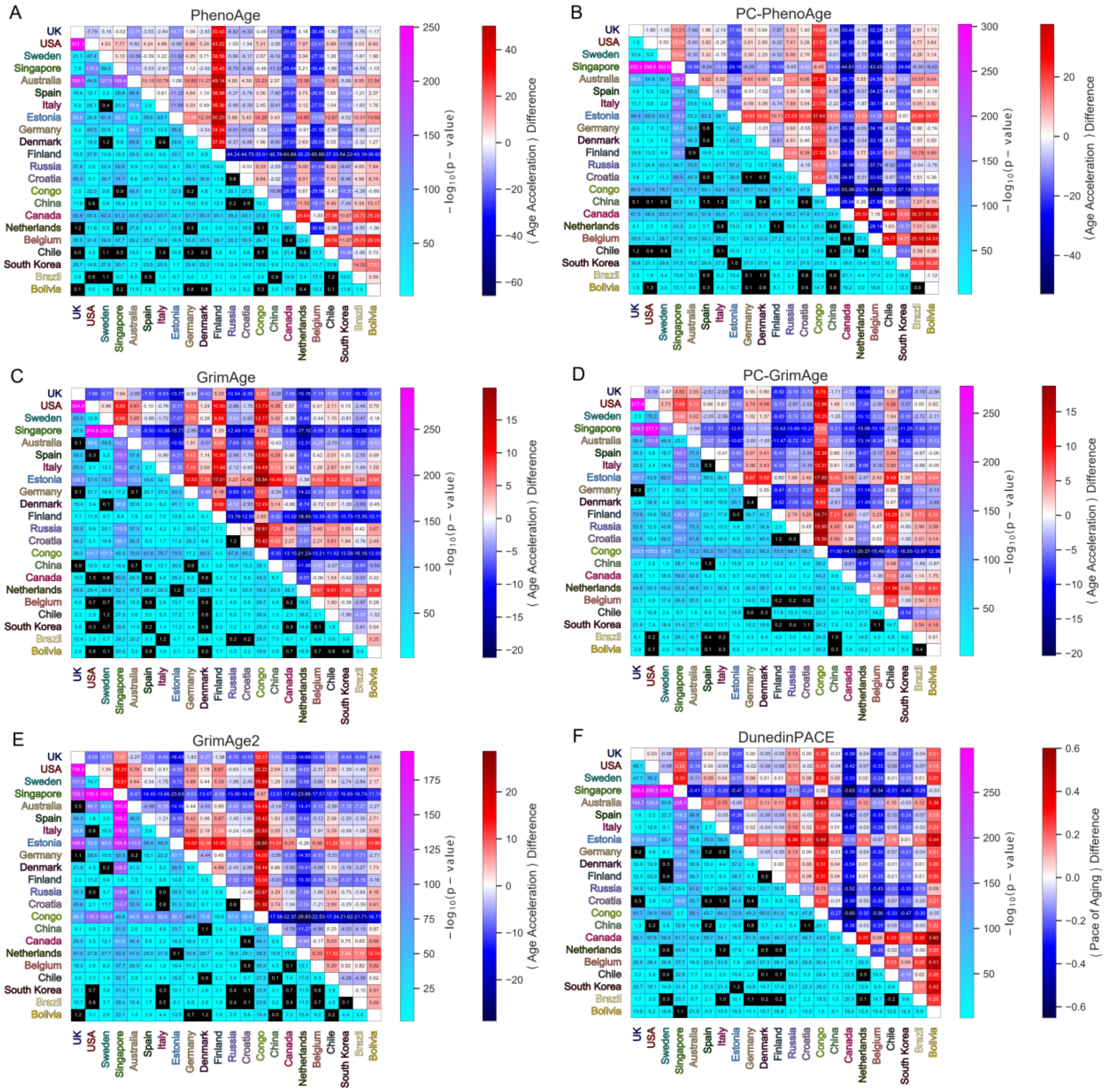


Supplementary Figure S4. Statistical significance of the difference in mean (A) DNAm PhenoAge, (B) PC-PhenoAge, (C) GrimAge, (D) PC-GrimAge, (E) GrimAge2, (F) DunedinPACE values between different countries for blood. Lower triangle: FDR-corrected pairwise Mann-Whitney U test p-value. Upper triangle: Difference between mean (A) DNAm PhenoAge, (B) PC-PhenoAge, (C) GrimAge, (D) PC-GrimAge, (E) GrimAge2, (F) DunedinPACE values (column minus row).

### 3. Tissues: countries and populations

In the main text, we discuss epigenetic age acceleration in different countries and populations for DNA methylation data from blood, as it is the biggest one. Epigenetic age acceleration for other tissues in different countries and populations will be considered below. Since we analyze tissues other than blood, we keep only pan-tissue epigenetic models - these are Horvath DNAmAge and SkinBlood DNAmAge with their PC-modifications.

##### 3.1. Buccal

The most numerous after blood is buccal methylation data. It shows quite a good diversity of representatives of different countries: Canada, UK, Netherlands, USA, Jamaica (Supplementary Figure S5A). The most samples are from Canada (almost 1000), while representatives of other countries are much less (Supplementary Figure S5B). The age distributions of the samples differ significantly - fairly wide age ranges for the USA and Jamaica (15-20 to 50-60 years), almost only children for Canada, UK and Netherlands. To investigate how pan-tissue epigenetic models work for different countries across age ranges, we calculated the moving average (within 5 years) for epigenetic age acceleration values as a function of the real age of the samples (Supplementary Figure S5C). The original versions of the epigenetic clock work rather well for buccal - the age acceleration values do not take extremely large positive or negative values for all samples in all age ranges. Interestingly, for PC-Horvath, the Canadian data tend to show a monotonically decreasing epigenetic acceleration with age. The Jamaican samples show positive age acceleration for all epigenetic estimates over almost the entire age range (except Horvath and PC-Horvath for samples older than 50 years). The average values of epigenetic age acceleration for different countries and different epigenetic models are plotted as a hierarchical clustering diagram (Supplementary Figure S5D). For Jamaica, all pan-tissue epigenetic models show positive age acceleration, with the highest value demonstrated by PC-SkinBloodAge. In the hierarchical clustering, Jamaica is separated from the two groups formed by Canada and the Netherlands (perhaps they are similar because they are mostly children there), UK and USA. If we consider hierarchical clustering for the epigenetic models, the original epigenetic estimates are very similar in terms of mean values of age acceleration, the PC-modifications differ from them and among themselves. We also, for each country, divided the entire data into females and males, and applied the Mann-Whitney U-test to these two groups, obtaining FDR-adjusted p-values. We then calculated the number of epigenetic clocks that have higher epigenetic age acceleration in males than females (blue bars in Supplementary Figure S5E), higher epigenetic age acceleration in females than males (red bars in Supplementary Figure S5E), and no statistical significance/one sex (gray bars in Supplementary Figure S5E). Males have higher epigenetic age acceleration than females in the USA. Females have a higher epigenetic age acceleration than males in Canada and Netherlands. In the remaining countries, there are either no statistical differences between males and females or the country is represented by only one sex.

All countries except the USA have representatives of only one population, so we consider the populations within the USA (White Americans, African Americans, Latinos) separately (Supplementary Figure S5F). White Americans and African Americans are about the same in quantity with similar age distributions (20-60 years), Latinos are very few - only 5 samples with a smaller age distribution (30-55 years). The moving average of epigenetic age acceleration with real age behaves similarly in all populations and epigenetic estimates (Supplementary Figure S5G): White Americans and African Americans have age acceleration around 0 over almost the entire age range, with overestimation for White Americans around 40-45 years and underestimation for White Americans around 50-55 years. Latinos have negative mean age acceleration for all estimates except PC-SkinBloodAge, but still they are not numerous enough to be sufficiently representative. The diagram of mean values of age acceleration by populations with hierarchical clustering (Supplementary Figure S5H) is consistent with the previous results - White Americans and African Americans are similar in all epigenetic models, Latinos show negative age acceleration (most likely due to dataset peculiarities). Males in White Americans and African Americans have higher epigenetic age acceleration than females (Supplementary Figure S5I).

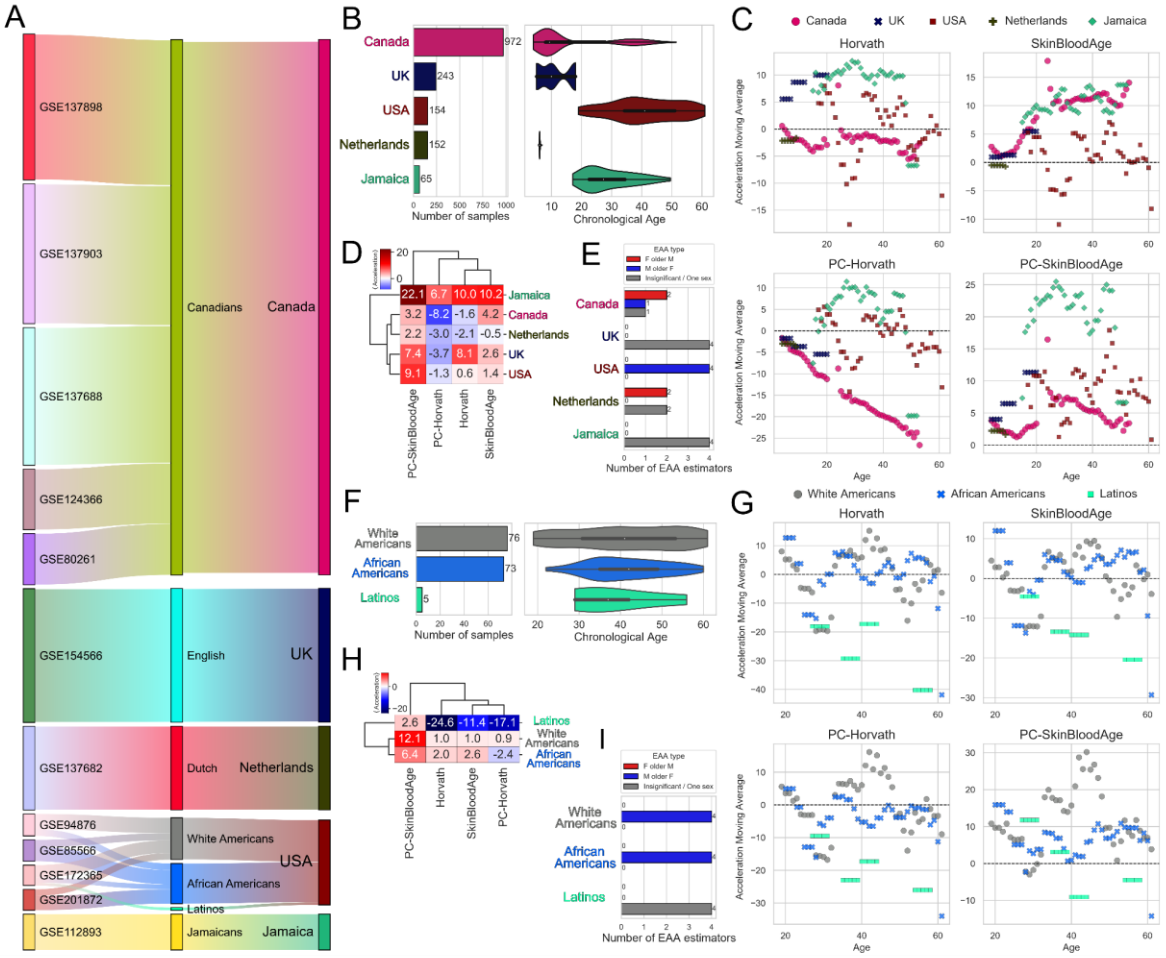


Supplementary Figure S5. Epigenetic age acceleration in different countries and populations in the USA in Buccal. (A) Sankey plot showing relations between datasets (GSEs, left), populations (center) and countries (right). (B) Number of samples from different countries (left) with corresponding age distributions (right). (C) Dependence of the moving average (within 5 years) of epigenetic age acceleration on real age for all countries for pan-tissue epigenetic models. (D) Diagram of the mean values of epigenetic age acceleration for countries for pan-tissue epigenetic clocks with hierarchical clustering. (E) Number of epigenetic metrics for each country for which: males have higher epigenetic age acceleration (blue), females have higher epigenetic age acceleration (red), no statistical significance/one sex (gray). Mann-Whitney U-test was applied for males and females for each tissue, with FDR-corrected resulting p-values. (F) Number of samples from different populations in the USA (left) with corresponding age distributions (right). (G) Dependence of the moving average (within 5 years) of epigenetic age acceleration on real age for all populations in the USA for pan-tissue epigenetic models. (H) Diagram of the mean values of epigenetic age acceleration for populations in the USA for pan-tissue epigenetic clocks with hierarchical clustering. (I) Number of epigenetic metrics for each population in the USA for which: males have higher epigenetic age acceleration (blue), females have higher epigenetic age acceleration (red), no statistical significance/one sex (gray). Mann-Whitney U-test was applied for males and females for each tissue, with FDR-corrected resulting p-values.

##### 3.2. Brain

The next in the number of samples is brain methylation data. It also shows quite a good diversity of representatives from different countries: USA, UK, Denmark, Canada, Australia, Germany (Supplementary Figure S6A). The largest number of samples is from the USA (more than 800), the number of samples from other countries is much smaller (Supplementary Figure S6B). The age distributions of the samples differ significantly - quite wide age ranges for USA and UK (15-90 years old), almost only elderly for Australia, Germany and Denmark (over 60 years). Let us consider the moving average (within 5 years) for epigenetic age acceleration values as a function of the real age of the samples (Supplementary Figure S6C). All clocks perform similarly for all samples - almost monotonically decreasing from positive and zero values for young ages to strongly negative values (over 30-40 years) for older ages. The average values of epigenetic age acceleration for different countries and different epigenetic models are plotted with hierarchical clustering (Supplementary Figure S6D). All epigenetic estimates for all countries show strongly negative values - this appears to be a specific feature of brain methylation. For all countries except Denmark, there are no statistically significant differences between males and females. In Denmark, males have a higher epigenetic age acceleration (Supplementary Figure S6E).

All countries except the USA have representatives of only one population, so we consider the populations within the USA (White Americans, African Americans) separately (Supplementary Figure S6F). White Americans and African Americans are about the same in number with a similar age distribution (10-80 years). The moving average (within 5 years) of epigenetic age acceleration with age behaves similarly in both populations and all epigenetic estimates (Supplementary Figure S6G): it decreases almost monotonically with age from zero to strongly negative values. The plot of mean values of age acceleration across populations with hierarchical clustering (Supplementary Figure S6H) is consistent with previous results - White Americans and African Americans are similar across all epigenetic models. No statistically significant difference was found between males and females (Supplementary Figure S6I).

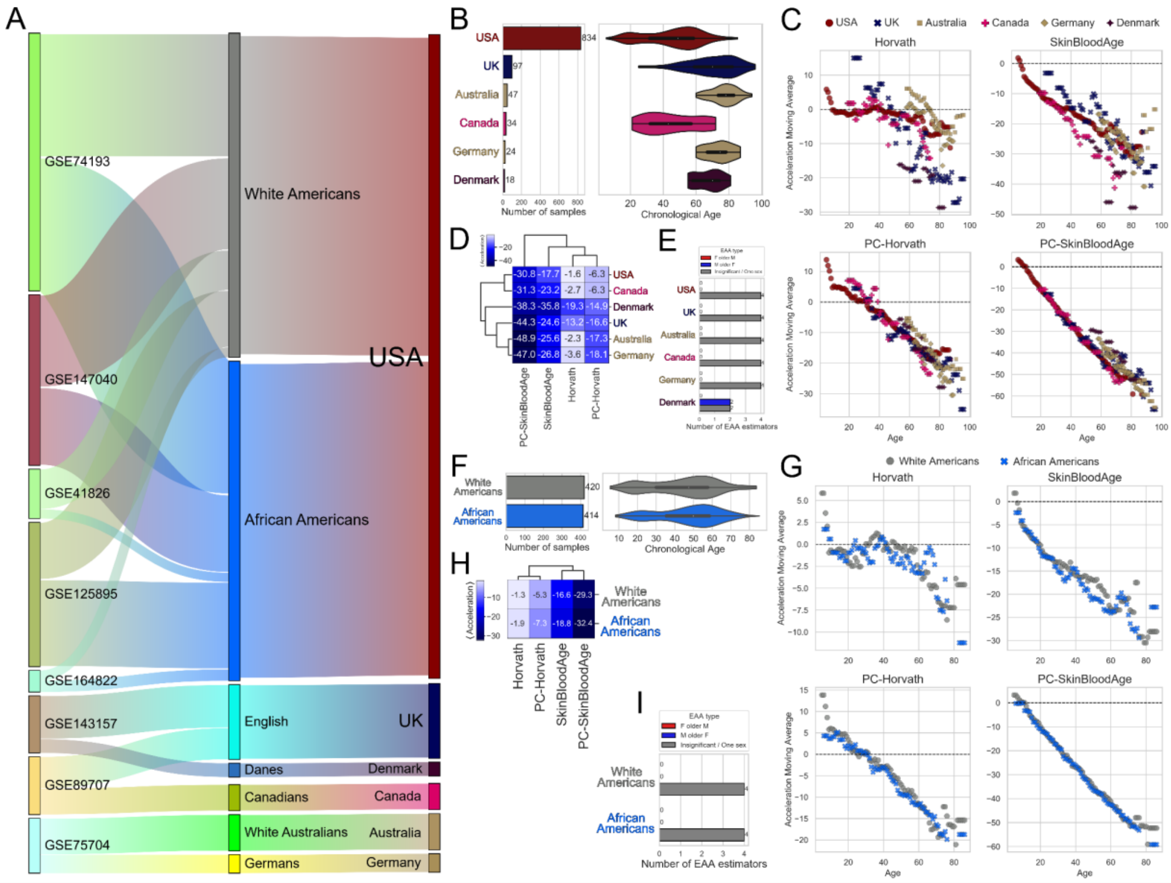


Supplementary Figure S6. Epigenetic age acceleration in different countries and populations in the USA in Brain. (A) Sankey plot showing relations between datasets (GSEs, left), populations (center) and countries (right). (B) Number of samples from different countries (left) with corresponding age distributions (right). (C) Dependence of the moving average (within 5 years) of epigenetic age acceleration on real age for all countries for pan-tissue epigenetic models. (D) Diagram of the mean values of epigenetic age acceleration for countries for pan-tissue epigenetic clocks with hierarchical clustering. (E) Number of epigenetic metrics for each country for which: males have higher epigenetic age acceleration (blue), females have higher epigenetic age acceleration (red), no statistical significance/one sex (gray). Mann-Whitney U-test was applied for males and females for each tissue, with FDR-corrected resulting p-values. (F) Number of samples from different populations in the USA (left) with corresponding age distributions (right). (G) Dependence of the moving average (within 5 years) of epigenetic age acceleration on real age for all populations in the USA for pan-tissue epigenetic models. (H) Diagram of the mean values of epigenetic age acceleration for populations in the USA for pan-tissue epigenetic clocks with hierarchical clustering. (I) Number of epigenetic metrics for each population in the USA for which: males have higher epigenetic age acceleration (blue), females have higher epigenetic age acceleration (red), no statistical significance/one sex (gray). Mann-Whitney U-test was applied for males and females for each tissue, with FDR-corrected resulting p-values.

##### 3.3. Colon

Next, let us consider the colon data. There are representatives of only three countries: USA, Poland, Ireland (Supplementary Figure S7A). The largest number of samples is from the USA (almost 500), the others are much less (Supplementary Figure S7B). The age distributions of the samples are different - bimodal distribution for the USA (5-20 years and 40-80 years), middle-aged and elderly for Poland and Denmark (50-80 years and 40-70 years, respectively). Let us consider the moving average (within 5 years) for epigenetic age acceleration values as a function of the real age of the samples (Supplementary Figure S7C). The clock performs similarly for all samples - increasing acceleration before age 40 and decreasing after age 40. The difference is that for SkinBloodAge and PC-SkinBloodAge the acceleration is always positive, while for Horvath and PC-Horvath the acceleration is positive until about age 50-55, then becomes negative. The mean values of epigenetic age acceleration for different countries and different epigenetic models are plotted with hierarchical clustering (Supplementary Figure S7D). SkinBloodAge and PC-SkinBloodAge show strongly positive values, while Horvath and PC-Horvath show small-value accelerations of different signs. For all countries, there are no statistically significant differences between males and females (Supplementary Figure S7E). All countries except the USA have representatives of only one population, so we consider the populations within the USA (White Americans, African Americans, Asian Americans) separately (Supplementary Figure S7F). African Americans are the most numerous, mostly 40-70 years; White Americans are smaller, with a wide age range (5-80 years); Asian Americans are very few, only 5 samples with a small age range (5-15 years). The moving average (within 5 years) of epigenetic age acceleration with age behaves similarly across all populations and epigenetic estimates (Supplementary Figure S7G) and follows the general trend displayed in Supplementary Figure S7C. The plot of mean age acceleration values across populations with hierarchical clustering (Supplementary Figure S7H) is consistent with previous results, with SkinBloodAge and PC-SkinBloodAge showing strongly positive values, and Horvath and PC-Horvath showing slight acceleration of different signs. No statistically significant difference was found between males and females (Supplementary Figure S7I).

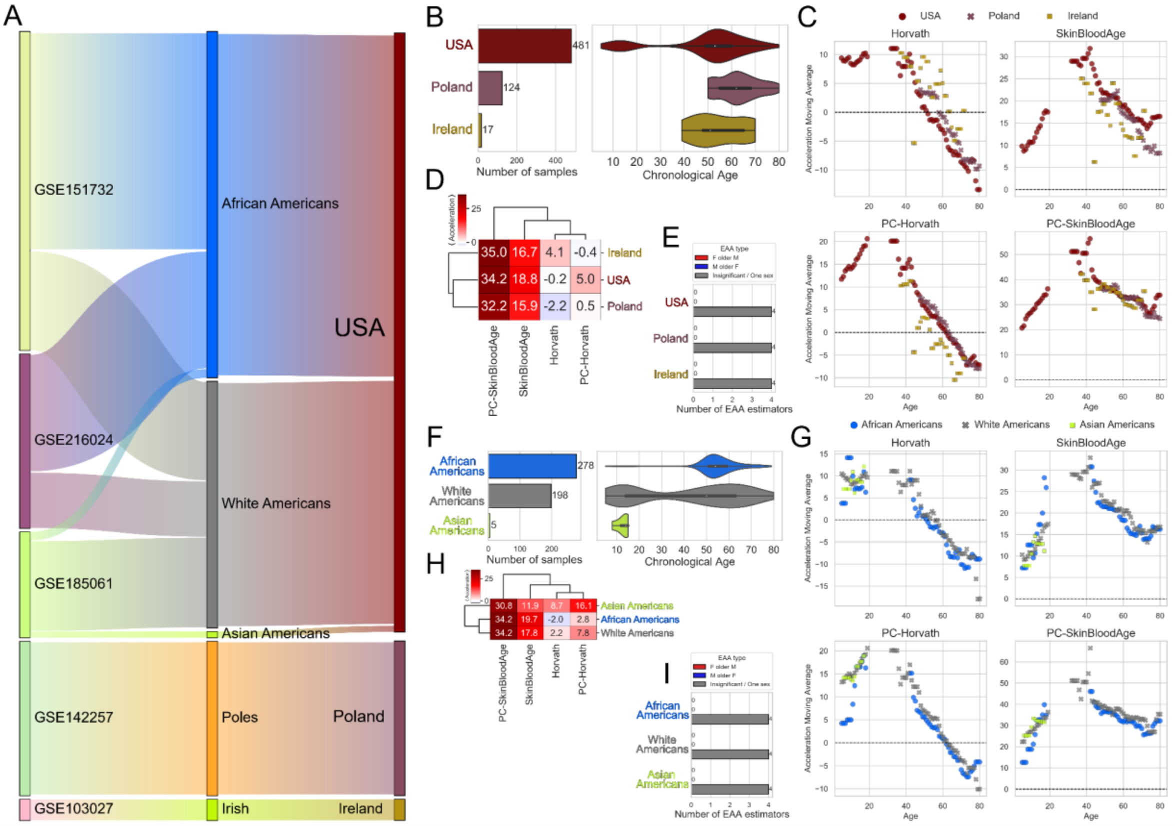


Supplementary Figure S7. Epigenetic age acceleration in different countries and populations in the USA in Colon. (A) Sankey plot showing relations between datasets (GSEs, left), populations (center)

and countries (right). (B) Number of samples from different countries (left) with corresponding age distributions (right). (C) Dependence of the moving average (within 5 years) of epigenetic age acceleration on real age for all countries for pan-tissue epigenetic models. (D) Diagram of the mean values of epigenetic age acceleration for countries for pan-tissue epigenetic clocks with hierarchical clustering. (E) Number of epigenetic metrics for each country for which: males have higher epigenetic age acceleration (blue), females have higher epigenetic age acceleration (red), no statistical significance/one sex (gray). Mann-Whitney U-test was applied for males and females for each tissue, with FDR-corrected resulting p-values. (F) Number of samples from different populations in the USA (left) with corresponding age distributions (right). (G) Dependence of the moving average (within 5 years) of epigenetic age acceleration on real age for all populations in the USA for pan-tissue epigenetic models. (H) Diagram of the mean values of epigenetic age acceleration for populations in the USA for pan-tissue epigenetic clocks with hierarchical clustering. (I) Number of epigenetic metrics for each population in the USA for which: males have higher epigenetic age acceleration (blue), females have higher epigenetic age acceleration (red), no statistical significance/one sex (gray). Mann-Whitney U-test was applied for males and females for each tissue, with FDR-corrected resulting p-values.

##### 3.4. Saliva

Next, we consider the saliva data. There are representatives of only three countries: USA, Netherlands, Belgium (Supplementary Figure S8A). The largest number of samples is from the USA (almost 200), the others are much smaller (Supplementary Figure S8B). The age distributions of the samples are different - wide distribution for the USA (20-90 years), children for the Netherlands and Belgium (around 10 years). Let us consider the moving average (within 5 years) for the epigenetic age acceleration values as a function of the real age of the samples (Supplementary Figure S8C). Both PC versions of the clocks perform similarly, with weakly negative age acceleration for children from the Netherlands and weakly positive age acceleration for children from Belgium. For the US, the PC clocks show a decrease in age acceleration with age. For children from the Netherlands and Belgium, the original versions of the clocks show weakly negative age acceleration. For the USA, the Horvath model shows decreasing age acceleration with age, the SkinBloodAge model always shows positive age acceleration with two maxima around 35 years and 60 years. The mean values of epigenetic age acceleration for different countries and different epigenetic models are plotted with hierarchical clustering (Supplementary Figure S8D). All epigenetic metrics show small values of mean age acceleration, hierarchical clustering unites the Netherlands with Belgium (indeed, both are children). For all countries except the USA, there are no statistically significant differences between males and females. In the USA, males have a higher epigenetic age acceleration (Supplementary Figure S8E).

All countries except the USA have representatives of only one population, so let's consider the populations within the USA (White Americans, Latinos, African Americans, Asian Americans) separately (Supplementary Figure S8F). White Americans are the most numerous, while other populations are less numerous. White Americans and Latinos have the widest age range (20-85 years), African Americans have a slightly narrower range (20-70 years), and Asian Americans have only two young adults (just over 20 years). The moving average (within 5 years) of epigenetic age acceleration with age behaves similarly in all populations (Supplementary Figure S8G) and follows the general trend shown in Supplementary Figure S8C. The plot of mean age acceleration across populations with hierarchical clustering

(Supplementary Figure S8H) shows that Horvath and PC-Horvath demonstrate negative mean acceleration for all populations, and SkinBloodAge and PC-SkinBloodAge demonstrate positive mean acceleration (except for PC-SkinBloodAge for Latinos, which is -0.4). For all populations except African Americans, there are no statistically significant differences between males and females. For African Americans, males have a higher epigenetic age acceleration (Supplementary Figure S8I).

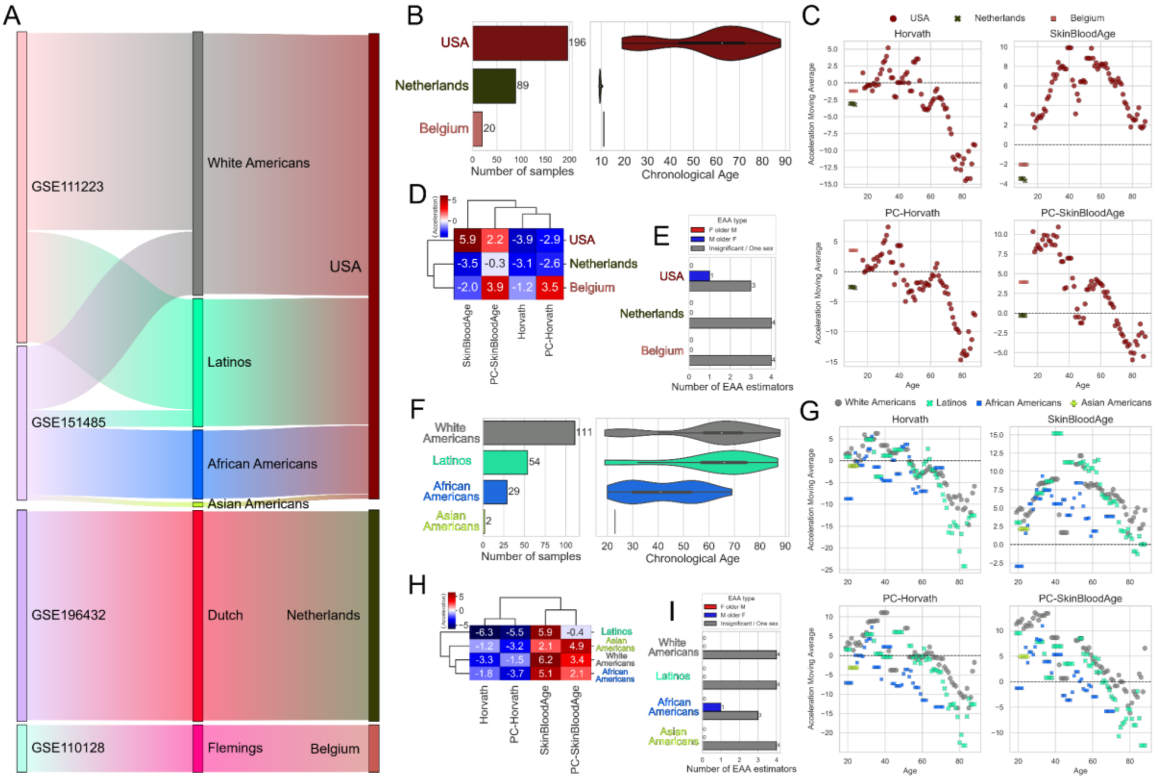


Supplementary Figure S8. Epigenetic age acceleration in different countries and populations in the USA in Saliva. (A) Sankey plot showing relations between datasets (GSEs, left), populations (center) and countries (right). (B) Number of samples from different countries (left) with corresponding age distributions (right). (C) Dependence of the moving average (within 5 years) of epigenetic age acceleration on real age for all countries for pan-tissue epigenetic models. (D) Diagram of the mean values of epigenetic age acceleration for countries for pan-tissue epigenetic clocks with hierarchical clustering. (E) Number of epigenetic metrics for each country for which: males have higher epigenetic age acceleration (blue), females have higher epigenetic age acceleration (red), no statistical significance/one sex (gray). Mann-Whitney U-test was applied for males and females for each tissue, with FDR-corrected resulting p-values. (F) Number of samples from different populations in the USA (left) with corresponding age distributions (right). (G) Dependence of the moving average (within 5 years) of epigenetic age acceleration on real age for all populations in the USA for pan-tissue epigenetic models. (H) Diagram of the mean values of epigenetic age acceleration for populations in the USA for pan-tissue epigenetic clocks with hierarchical clustering. (I) Number of epigenetic metrics for each population in the USA for which: males have higher epigenetic age acceleration (blue), females have higher epigenetic age acceleration (red), no statistical significance/one sex (gray). Mann-Whitney U-test was applied for males and females for each tissue, with FDR-corrected resulting p-values.

##### 3.5. Lung

Next, let's analyze lung data. There are representatives of only one country, the USA, but there are 4 populations - White Americans, African Americans, Latinos, Asian Americans (Supplementary Figure S9A). White Americans are the most numerous (over 150), while the other populations are smaller (Supplementary Figure S9B). White Americans and African Americans have the widest age range (15-85 and 15-75 years, respectively), there are 8 Latinos (50-65 years), and there is only one Asian American (just over 70 years). We consider the moving average (within 5 years) for the epigenetic age acceleration values as a function of the real age of the samples (Supplementary Figure S9C). Both PC versions of the clocks work in a similar way - decreasing age acceleration with age. The Horvath model shows decreasing age acceleration with age, the SkinBloodAge model almost always shows positive age acceleration with a maximum around age 60. The mean values of epigenetic age acceleration for different populations and different epigenetic models are plotted with hierarchical clustering (Supplementary Figure S9D). PC-SkinBloodAge shows negative age acceleration for all studied populations, all other epigenetic estimates show positive age acceleration. No statistically significant difference was found between males and females (Supplementary Figure S9E).

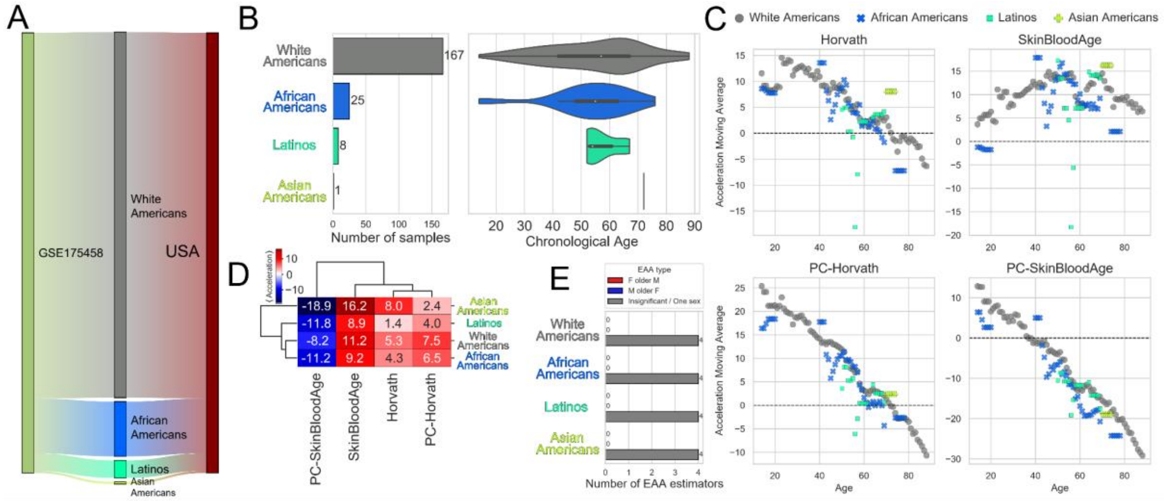


Supplementary Figure S9. Epigenetic age acceleration in different populations in the USA in Lung. (A) Sankey plot showing relations between datasets (GSEs, left), populations (center) and countries (right). (B) Number of samples from different populations (left) with corresponding age distributions (right). (C) Dependence of the moving average (within 5 years) of epigenetic age acceleration on real age for all populations for pan-tissue epigenetic models. (D) Diagram of the mean values of epigenetic age acceleration for populations for pan-tissue epigenetic clocks with hierarchical clustering. (E) Number of epigenetic metrics for each population for which: males have higher epigenetic age acceleration (blue), females have higher epigenetic age acceleration (red), no statistical significance/one sex (gray). Mann-Whitney U-test was applied for males and females for each tissue, with FDR-corrected resulting p-values.

##### 3.6. Breast

Let us consider the breast data further. There are members of only one country, USA, but there are 2 populations: White Americans, African Americans (Supplementary Figure S10A). White Americans are more numerous than African Americans (Supplementary Figure S10B). White

Americans have a wider age range (15-75 years) than African Americans (20-55 years). We consider the moving average (within 5 years) for epigenetic age acceleration values as a function of the real age of the samples (Supplementary Figure S10C). All epigenetic clocks work in a similar way - decreasing age acceleration with age, with always positive values. The mean values of epigenetic age acceleration for different populations and different epigenetic models are plotted with hierarchical clustering (Supplementary Figure S10D). All epigenetic estimates show positive mean age acceleration for all studied populations. No statistically significant difference was found between males and females (Supplementary Figure S10E).

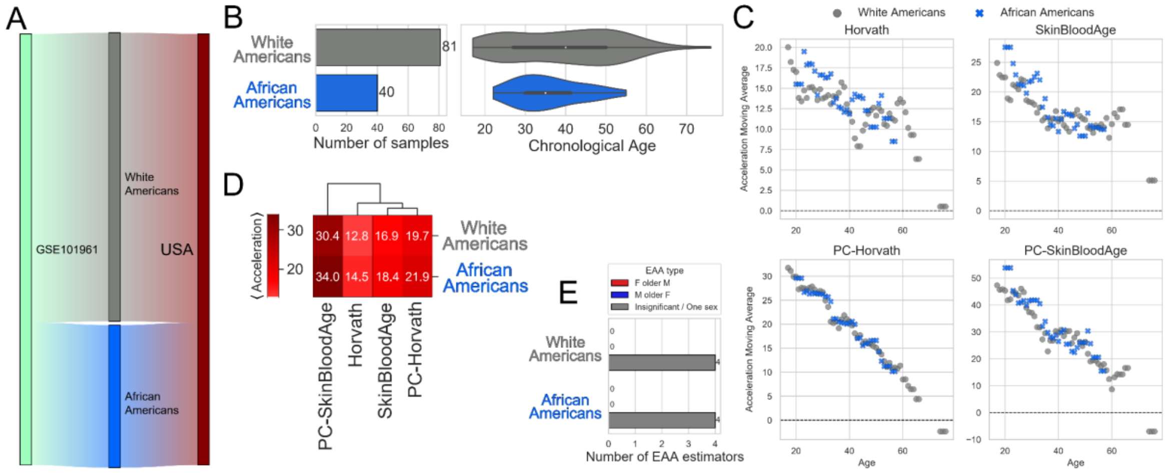


Supplementary Figure S10. Epigenetic age acceleration in different populations in the USA in Breast. (A) Sankey plot showing relations between datasets (GSEs, left), populations (center) and countries (right). (B) Number of samples from different populations (left) with corresponding age distributions (right). (C) Dependence of the moving average (within 5 years) of epigenetic age acceleration on real age for all populations for pan-tissue epigenetic models. (D) Diagram of the mean values of epigenetic age acceleration for populations for pan-tissue epigenetic clocks with hierarchical clustering. (E) Number of epigenetic metrics for each population for which: males have higher epigenetic age acceleration (blue), females have higher epigenetic age acceleration (red), no statistical significance/one sex (gray). Mann-Whitney U-test was applied for males and females for each tissue, with FDR-corrected resulting p-values.

##### 3.7. Other tissues

For each of the tissues, epidermis (Supplementary Figure S11A), muscle (Supplementary Figure S11B) and liver (Supplementary Figure S11C), there is data from only one country. Each of these countries has members of only one population. Therefore, the epigenetic age acceleration analysis for these tissues is fully presented in the main text of the paper, Figure 2.

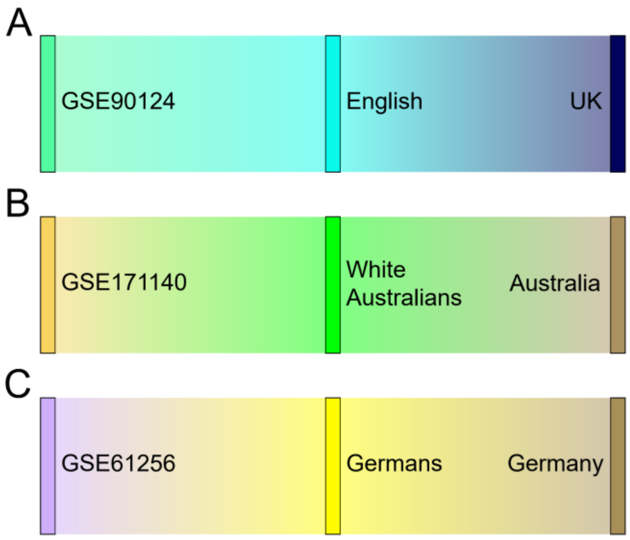


Supplementary Figure S11. Sankey plot showing relations between datasets (GSEs, left), populations (center) and countries (right) for (A) Epidermis, (B) Muscle, (C) Liver.
